# Resolving the transcriptional transitions associated with oligodendrocyte generation from adult neural stem cells by single cell sequencing

**DOI:** 10.1101/2020.12.18.423285

**Authors:** Kasum Azim, Filippo Calzolari, Martina Cantone, Rainer Akkermann, Julio Vera, Hans-Peter Hartung, Onur Basak, Arthur Morgan Butt, Patrick Küry

## Abstract

The subventricular zone (SVZ) is the largest neurogenic niche in the adult forebrain. Notably, neural stem cells (NSCs) of the SVZ generate not only neurons, but also oligodendrocytes, the myelin-forming cells of the central nervous system. Transcriptomic studies have provided detailed knowledge of the molecular events that regulate neurogenesis, but little is understood about adult oligodendrogenesis from SVZ-NSCs. To address this, we performed in-depth single-cell transcriptomic analyses to resolve the major differences in neuronal and oligodendroglial lineages derived from the adult SVZ. A hallmark of adult oligodendrogenesis was the stage-specific expression of transcriptional modulators that regulate developmental oligodendrogenesis. Notably, divergence of the oligodendroglial lineage was distinguished by Wnt-Notch and angiogenesis-related signaling, whereas G-protein-coupled receptor signaling pathways were the major signature observed in the neurogenic lineage. Moreover, in-depth gene regulatory network analysis identified key stage-specific master regulators of the oligodendrocyte lineage and revealed new mechanisms by which signaling pathways interact with transcriptional networks to control lineage progression. Our work provides an integrated view of the multi-step differentiation process leading from NSCs to mature oligodendrocytes, by linking environmental signals to known and novel transcriptional mechanisms orchestrating oligodendrogenesis.

**Main points:**

- Distinct adult NSC populations giving rise to either oligodendrocytes or neurons can be identified by the expression of transcription factors.
- Gene regulatory control of oligodendrogenesis is a major fate-determinant for their generation.

## INTRODUCTION

Oligodendrocytes are the myelin-forming cells of the central nervous system (CNS), and provide insulation for rapid axonal conduction and support to axons. In the forebrain, most oligodendrocytes are generated shortly after birth from neural stem cells (NSCs) of the subventricular zone (SVZ) (Azim, Berninger, & Raineteau, 2016; Kessaris et al., 2006), which are also responsible for neurogenesis (Figueres-Onate, Sanchez-Villalon, Sanchez-Gonzalez, & Lopez-Mascaraque, 2019), and this activity is retained well into late adulthood (Fuentealba et al., 2015). The sequence of oligodendroglial differentiation is relatively similar in postnatal and adult SVZ (Azim et al., 2016; El Waly, Macchi, Cayre, & Durbec, 2014), whereby NSCs first generate transiently amplifying progenitors (TAPs), characterized by elevated expression of the transcription factor *Ascl1* (Nakatani et al., 2013), followed by progressive differentiation into oligodendrocyte precursor cells (OPCs) and maturation into myelinating oligodendrocytes (Azim et al., 2016; El Waly et al., 2014). In rodents, approximately 5% of all cells derived from the adult SVZ are oligodendrocyte lineage cells (Capilla-Gonzalez, Cebrian-Silla, Guerrero-Cazares, Garcia-Verdugo, & Quinones-Hinojosa, 2013; El Waly et al., 2014; Menn et al., 2006).

Fate-mapping studies using Cre drivers under the control of regulatory regions from pallial and subpallial transcription factors demonstrated that molecularly distinct SVZ microdomains derive from their embryonic counterparts (reviewed in (Azim et al., 2016)). The embryonic septum, the lateral ganglionic eminences, and cortex contain NSCs that populate the medial (i.e. septal), ventral (i.e. striatal; also called lateral) and dorsal (i.e. cortical) aspects of the adult SVZ, respectively (Azim et al., 2016; Fiorelli, Azim, Fischer, & Raineteau, 2015). Localized labeling of NSCs showed that regionally segregated NSCs are robustly committed to generating distinct cell subtypes (Menn et al., 2006; Merkle et al., 2014; Merkle, Mirzadeh, & Alvarez-Buylla, 2007), and that environmental signals impinge on intrinsic fate control mechanisms to modulate specific aspects of NSC and progeny behavior.

The OPC and oligodendrocyte transcriptional programs in both adult and postnatal contexts have been comprehensively dissected using single cell sequencing technologies (Marques et al., 2018; Marques et al., 2016; Zeisel et al., 2015). A recent study revealed regional differences in lineage output from the SVZ with its ventral microdomain exhibiting predominantly pro-neuronal-, and the medial wall exhibiting pro-oligodendroglial hallmarks (Mizrak et al., 2019), in line with previous observations using fate-mapping and transcriptomic profiling methods demonstrating that ventral SVZ-NSCs during postnatal development (Azim, Fischer, et al., 2014; Azim et al., 2015; Zweifel et al., 2018) and in the adult (Azim, Raineteau, & Butt, 2012; Dulken, Leeman, Boutet, Hebestreit, & Brunet, 2017; Llorens-Bobadilla et al., 2015; Felipe Ortega et al., 2013) are biased towards olfactory bulb neurogenesis. The generation of oligodendrocytes from the SVZ dorsal wall during early postnatal development is, however, considered as the default source for oligodendrocyte generation (Azim et al., 2017; Azim, Fischer, et al., 2014; Azim, Rivera, Raineteau, & Butt, 2014; Kessaris et al., 2006). Likewise, the adult dorsal SVZ serves as an additional microdomain for their generation (Azim et al., 2017; Azim et al., 2012; Crawford, Tripathi, Richardson, & Franklin, 2016; Felipe Ortega et al., 2013). In contrast to the number of high throughput sequencing studies defining the molecular events underlying neuronal generation from adult NSC subpopulations (Marcy & Raineteau, 2019), far less is known about the regulatory context of adult SVZ-dependent oligodendrogenesis. This is partly due to technical limitations in isolating relevant microdomains, e.g. typical SVZ whole-mount preparations are devoid of dorsal SVZ tissue (Mirzadeh, Doetsch, Sawamoto, Wichterle, & Alvarez-Buylla, 2010). To overcome such restrictions, a cruder approach was used by capturing both dorsal and lateral walls (Basak et al., 2018), which enabled us to sequence sufficient numbers of single cells and to examine lineage-specific cells in the adult SVZ. Here, we describe the molecular events that regulate oligodendrogenesis from the adult SVZ and unravel mechanisms by which transcription factor networks and signaling pathways interact to drive specification in the early oligodendrocyte lineage.

## MATERIALS AND METHODS

### Determining the lineage memberships of SVZ-derived single NSCs and TAPs

The procedures for transgenic and FACs strategies used for capturing single cells, mapping, pre-processing and RACEID are applied as described previously (Basak et al., 2018). The following reporter mice were used: quiescent(q)NSCs, few active(a) and primed(p)NSCs were taken from Troy:::GFP mice; TAPs and neuroblasts were harvested from Ki67:::RFP mice, and qNSCs (including a few pNSCs and aNSCs) were extracted from Ki67:::creRT2:::Tom2 mice. FACs analysis (Figure S1a) allowed identification/isolation of GLAST+/EGF-qNSCs, GLAST+/EGF+ aNSCs, GLAST-EGF+ TAPs and O4+ oligodendrocytes. Parameters for cell clusters identification were adapted as done previously (Basak et al., 2018) to obtain SVZ-lineage specific clusters.

We predicted that the transcriptional hallmarks of oligodendrogenesis and neurogenesis from the postnatal SVZ microdomains are retained to a certain extent into adulthood. Bulk datasets of our previous study (Azim et al., 2015) were used for identifying transcriptional signatures of the entire oligodendrocyte lineage by comparing dorsal NSCs, dorsal TAPs and oligodendrocytes and for ventral SVZ-derived neurogenesis. Transcription factors used as markers for lineage identification in adult single cell were (i) highly expressed during early postnatal development and (ii) uniquely expressed in either oligodendrocyte or neuronal lineages. These included among other genes *Notch2, Epas1, Foxo1,* and *Esrrg2*. A first characterization was performed with RACEID to identify rarer cell populations in single cell sequencing (Grun et al., 2015), then PartekFlow® (http://www.partek.com/partek-flow/) was used to assign single cells to defined SVZ lineages, including the rarer ones from RACEID. Visual inspection of t-Distributed Stochastic Neighbor Embedding (tSNE) plots displaying transcript levels for selected lineage-specific markers was used to sub-cluster cells found in close proximity in the plots. Figure S1b contains examples of some of the markers used in this oligodendrocyte lineage detection step.

Transcription factor markers obtained from bulk datasets were used to identify single cells of the oligodendroglial lineage (Azim et al., 2015), aside from OPCs and mature oligodendrocytes within subpopulations of NSCs/TAPs (Figure S1b). The soundness of our lineage membership assessment was further evaluated by comparing the gene expression profile of the remaining cells, expected to belong to the larger neurogenic population, with bulk datasets of adult qNSC/aNSC/TAP/NB populations. Adult TAPs and neuroblasts we described in a recent study (Azim et al., 2018).

PartekFlow was used to generate heat PCA plots describing pallial and subpallial marker expression by cells in the early stages of the oligodendrocyte lineage (qNSCI, qNSCII and pNSC). Gene Specific Analysis algorithm from PartekFlow was used with default parameters for acquiring gene lists defining the distinct cell lineages. Lineage-specificities were considered statistically relevant if the corresponding *p*-values were below 0.05. An additional analysis for determining the expression signatures of common genes (shared between oligodendroglial and neuronal lineages) was performed by identifying cluster-enriched genes for the first 5 stages of differentiation (qNSCI-to-TAP) without separation of lineage-specific clones. The above steps allowed the definition of a total of 13 clusters: qNSCI, qNSCII, pNSC, aNSC, TAP, neuroblasts for the neuronal lineage and OLqNSCI, OLqNSCII, OLpNSC, OLaNSC, OLTAP, OPC and mature oligodendrocytes for oligodendroglial lineage.

Differential expression analysis was performed using datasets mentioned above by applying methods as described in (Azim et al., 2018) in addition to the cluster enriched transcripts using the Seurat V3 package as described in the next paragraph. Genes were filtered to select for transcription factors or transcriptional cues identified with PartekFlow. Macro-groups accounting for distinct differentiation stages were characterized as follows: (i) early oligodendrocyte group and the neurogenic group were defined by their expression of transcription factors expressed in single cells of (OL)qNSCI, (OL)qNSCII and (OL)pNSC; (ii) mid group for either lineage included transcription factors expressed in aNSCs and TAPs. The same datasets were used to identify enriched transcription factors when specific differentiation stages were compared. Precisely, the following stages were examined as follows: (1) dorsal NSCs vs dorsal TAPs, OPCs, immature oligodendrocytes and mature oligodendrocytes; (2) dorsal TAPs vs dorsal NSCs, OPCs, immature oligodendrocytes and mature oligodendrocytes; (3) vNSCs vs vTAPs, TAPs and neuroblasts; (4) vTAPs vs vNSCs, TAPs and neuroblasts; (5) qNSCs vs aNSCs, TAPs and neuroblasts; (6) aNSCs vs qNSCs, TAPs and neuroblasts. Statistical analysis was performed using ANOVA for determining the differentially expressed (DE) transcription factors. Comparison of adult transcription factor expression in postnatal NSCs was summarized by assigning each transcription factor to a category: “1” indicates upregulated and highly expressed transcription factor, “-1” downregulated transcription factors, while “0” represents the remaining ones.

### Clustering and visualization of single cell data using the Seurat R package

The R package Seurat V3 was used to process single cell transcriptomic data and elucidate their heterogeneity (https://satijalab.org/seurat/vignettes.html (Butler, Hoffman, Smibert, Papalexi, & Satija, 2018)). Single cells were assigned a unique identifier, then the default pipeline for analysis was performed: data were log-normalized and cells containing mitochondrial genes with a *p*-value greater than 0.05 were filtered out. The obtained matrix was scaled so that JackStraw function could be applied for comparing the *p*-values distribution for all principal components. Using the genes displaying higher variation among the 13 clusters identified with PartekFlow, the Seurat function FindVariableFeatures determined 7 oligodendroglial and 6 neuronal clusters that were employed for the downstream pseudotime analysis. Graphical representations were created using Seurat package. Differential gene expression analysis was performed with FindAllMarkers function applying Wilcoxon rank sum test to find transcriptional markers of the 13 clusters. Of note, several statistical tests were implemented to ensure the reliability of the set of most relevant genes. Aside from DESeq2, differentially expressed genes across these methods were comparably similar (further detailed in https://satijalab.org/seurat/v3.0/de_vignette.html).

### FateID cellular trajectory construction

The genes specific for the 13 clusters were processed for assessing lineage biases by using FateID package for R environment. The goal is to resolve early lineage priming via iterative random forest classification (Herman, Sagar, & Grun, 2018). The default settings of the package were applied. In order to determine the relationship between terminal differentiation states and naïve stem/progenitor states, mature oligodendrocytes were equated as the heterotypic outlier of the neurogenic lineage. The variable genes in the neurogenic clusters and mature oligodendrocytes were used as a subset expression data frame. The top 10 genes belonging to each cluster as determined by the above Seurat analysis were used as learning set for the classification method hence defining the v matrix, while neuroblast and mature oligodendrocyte clusters were indicated as target set. Using fateBias function, a few iterations were operated by comparing consecutive stages of differentiation by setting the training set as the previous stage and the test set as the following one. For example, the defined NSC/TAP pool of the oligodendroglial lineage (including neuroblasts) was used as training set and mature oligodendrocytes, OPCs and neuroblasts were classified as the test set. Differentiation trajectories were computed using the principal curve function “prc”. The dimensionality reduction coordinates of the analyses of the cells and differentiation trajectories (data contained in the “dr” and “pr” lists) were exported and plotted using the ggplot2 R package for improved graphical representation.

### Cell cycle regression

NSCs/TAPs/neuroblasts, quiescent or activated, were further investigated to determine their position within the cell cycle, using CellCycleScoring function of Seurat package. Full details are available at https://satijalab.org/seurat/v3.0/cell_cycle_vignette.html (Butler et al., 2018). Seurat estimates the mean expression of the marker genes from 43 S-phase and 54 G2/M-phase thus generating a uniform rank for S- and G2/M-phase for each cell, which results in the assignment of each single cell to a particular phase. The score was gauged by deducting the mean expression of the 10 nearest neighbor-cells in dimensionality reduction space computed by Seurat. Cell cycle scorings of cell populations were plotted according to their pseudotime trajectory coordinates as previously calculated. The data are visualized as cell cycle marker signal distribution in ridge plots (Figure 3). The aNSCs and TAPs of both lineages were further classified according to the different phases of the cell cycle, therefore leading to the identification of 15 distinct clusters (while the initial aNSC and TAP clusters were replaced). Seurat was run generating cluster-enriched expression profiles and those of the mid stages were used for further GO analysis as described below.

**Figure 1.**
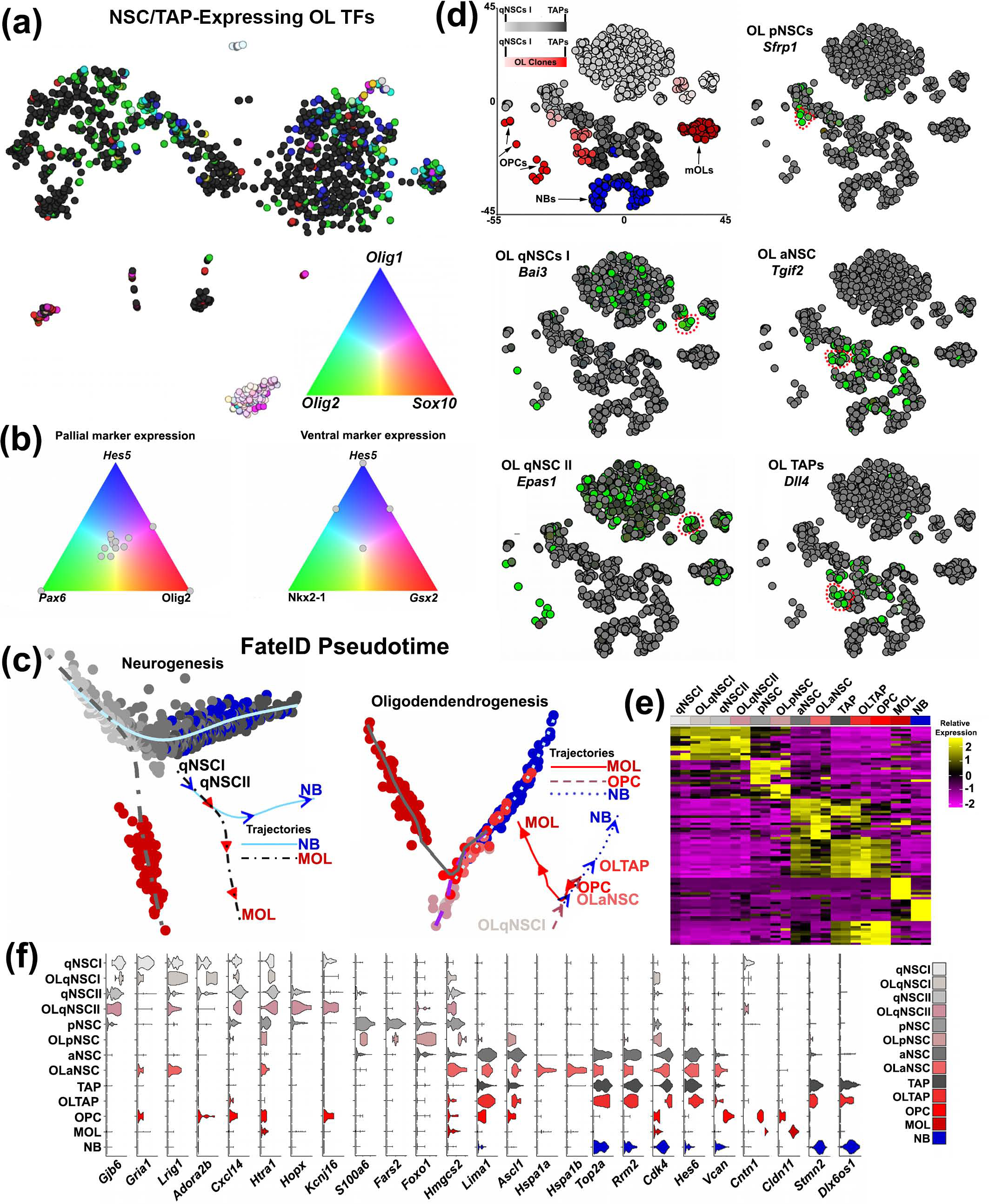
Identification of lineage-specific NSCs/NPs. (a) tSNE plot of 1200 single cells (including microglia, choroid plexus, neurons and vascular cells; see Figure S1) expressing the transcription factors *Olig1*, *Olig2* and *Sox10*. PCA color plot of transcript expression for determining cells with elevated expression of core oligodendrocyte lineage markers. (b) Selected early oligodendrocyte lineage cell expression of pallial marker *Pax6* and subpallial markers *Nkx2.1* and *Gsx2* together with the NSC marker *Hes5* for determining a potential dorsal origin of oligodendrocyte lineage. (c) FateID pseudotime demonstrates that the mature oligodendrocyte (MOL) transcriptional trajectory branches off only at the very earliest stages in the neuronal lineage, whereas among identified oligodendrocyte cells mature oligodendrocytes branch via OPCs that are in close proximity to OLTAPs and OLaNSCs. (d) tSNE plots of proneuronal and proolidendroglial lineage cells and selected transcript expression in the early- and mid-stages of the oligodendrocyte lineage. e Heatmap of the 13 analyzed clusters illustrating the highly enriched stage-specific signatures. Stage color legend is presented in f. (f) Violin plots of selected marker expression across the 13 clusters further demonstrating markers specific/enriched in stage and lineage. Abbreviations: aNSC: activated neural stem cell; MOL: mature oligodendrocyte; NB: neuroblast; NSC: neural stem cell; OLaNSC: oligodendroglial activated neural stem cell; OLpNSC: oligodendroglial primed neural stem cell; OLqNSCI: oligodendroglial quiescent neural stem cell (subtype I); OLqNSCI: oligodendroglial quiescent neural stem cell (subtype II); OLTAP: oligodendroglial transiently amplifying progenitor; OPC: oligodendrocyte precursor cell; pNSC: primed neural stem cell; qNSCI: quiescent neural stem cell (subtype I); qNSCII: quiescent neural stem cell (subtype II).

**Figure 2.**
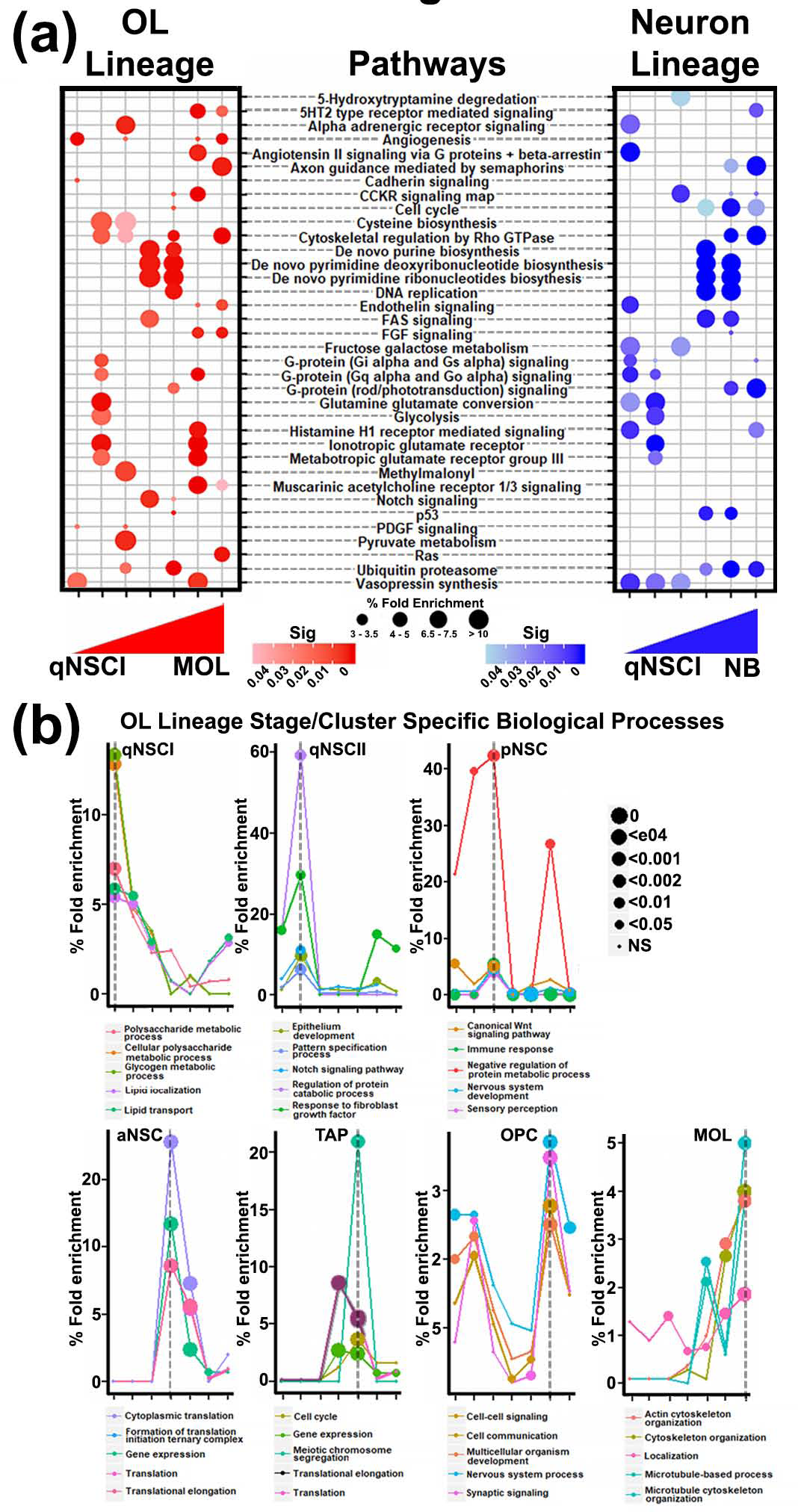
Pathway and mechanistic analysis of the oligodendrocyte lineage. (a) Dot plot of the most highly enriched/significant (“Sig” in plot) signaling pathways, organized alphabetically from top to bottom in the 2 lineages. (b) A time course of the top 5 most enriched biological processes in each of the 7 clusters in the oligodendrocyte lineage. Point size in plots reflects significance. Sig = significance (*p-*value). Abbreviations: aNSC: activated neural stem cell; MOL: mature oligodendrocyte; NB: neuroblast; NSC: neural stem cell; OPC: oligodendrocyte precursor cell; pNSC: primed neural stem cell; qNSCI: quiescent neural stem cell (subtype I); qNSCII: quiescent neural stem cell (subtype II).

**Figure 3.**
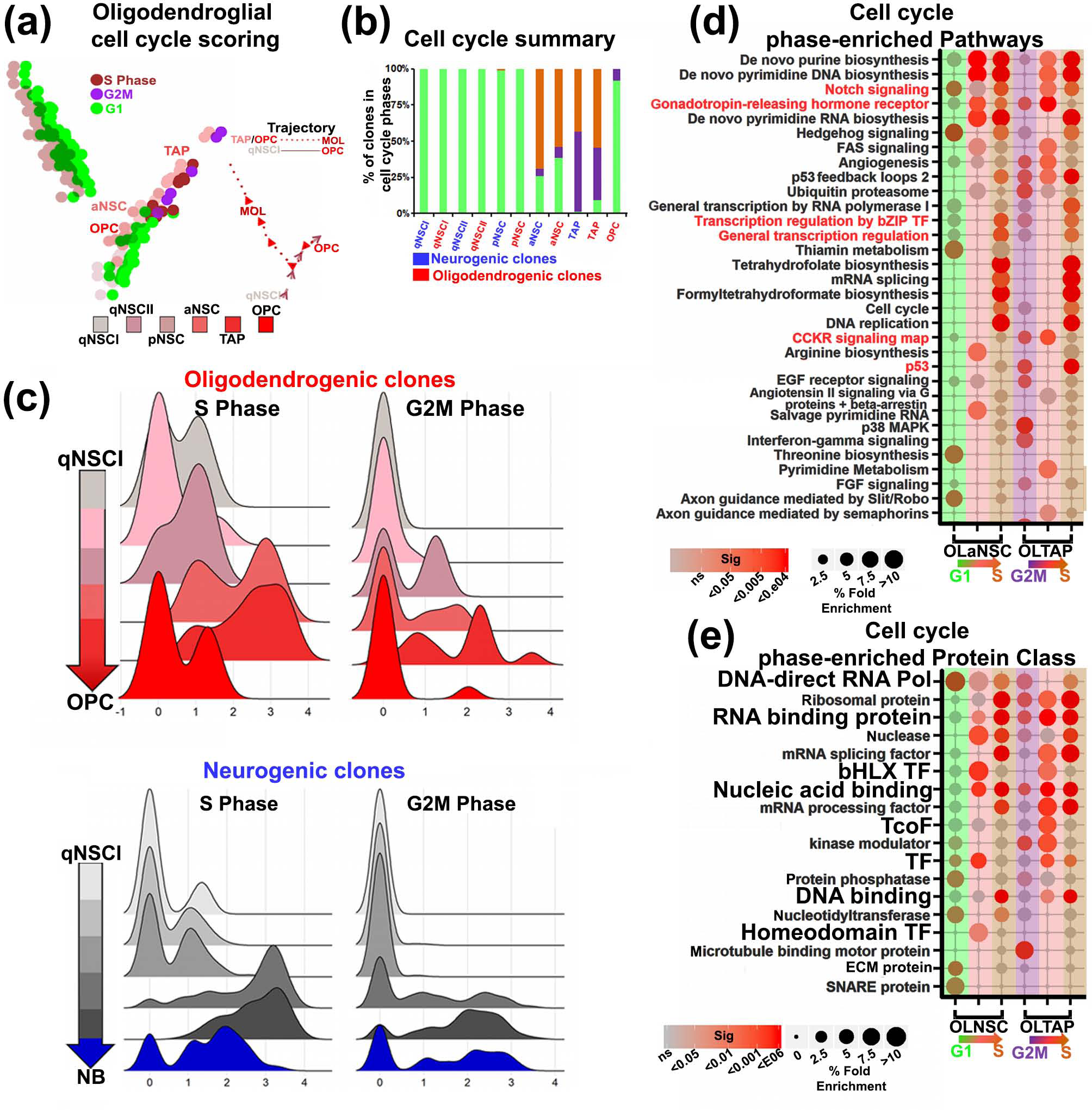
Cell cycle heterogeneity of oligodendroglial and neuronal lineage cells. (a) Oligodendrocyte and neuronal lineage cells scored by Seurat for cell cycle regression using canonical markers. Oligodendrocyte lineage cells are plotted according to their pseudotime trajectory and overlaid displayed the phase of cycle of each cell. (b) A summary displaying the percentage of cells in the 3 phases of cell cycle and the dynamics of mid stages of either lineage shuttling between states. (c) Ridge plot of oligodendrocyte and neuronal lineage cells illustrating the distribution of signal in the S or G2M cell cycle phase. (d) The mid stages of oligodendroglial (light red) and neuronal lineage (light blue) expanded for S (*Pcna, Mcm6*) and G2M (*Top2a, Mki67*) marker signal distribution in ridge plots. (e,f) Dot plots show pathway and protein class identification of the mid stages in the oligodendrocyte lineage. Green, light red and brown in OLaNSC labels quiescent, common to both stages and S phase, respectively; light purple, light red and light brown label OLTAPs in G2M, common to both and S phase, respectively. Terms are ranked according to their highest significance (Sig; *p-*value) from top to bottom. Pathways in red font are examined for revealing their signaling-to-transcriptional networks in Figure 7. Abbreviations: aNSC: activated neural stem cell; MOL: mature oligodendrocyte; NB: neuroblast; NSC: neural stem cell; OLaNSC: oligodendroglial activated neural stem cell; OLpNSC: oligodendroglial primed neural stem cell; OLqNSCI: oligodendroglial quiescent neural stem cell (subtype I); OLqNSCI: oligodendroglial quiescent neural stem cell (subtype II); OLTAP: oligodendroglial transiently amplifying progenitor; OPC: oligodendrocyte precursor cell; pNSC: primed neural stem cell; qNSCI: quiescent neural stem cell (subtype I); qNSCII: quiescent neural stem cell (subtype); TF: transcription factor.

### Gene ontology and pathway annotation of cell clusters

Gene lists for the original 13 clusters and the additional 4 ones (Figure 3e) were used for overrepresentation analysis using Protein ANalysis THrough Evolutionary Relationships (PANTHER) analysis tools ((http://www.pantherdb.org) (Mi, Poudel, Muruganujan, Casagrande, & Thomas, 2016)). Comparing the overrepresented pathways of the oligodendroglial and neuronal clusters, PANTHER pathways function was used for pointing out the mechanisms that differ or are shared between stages within a lineage or between the same stage in oligodendroglial and neuronal lineages. In this analysis, a minimum of 4 pathways for each cluster was retained if the corresponding *p*-value was below 0.05. TAP clusters or the later stage OPCs/mature oligodendrocytes contained a greater number of highly expressed genes, and so the top 10 most significantly expressed genes found with Seurat were used. Identification of genes that are unique to the oligodendrocyte lineage, the neuronal lineage and the shared ones was performed using the R Package Venn Diagram and the online tool http://www.biovenn.nl/index.php. Pathway and Protein Class lists obtained from PANTHER were downloaded for the analyses described above and presented as dot plots using the ggplot2 R package. Similarly, the time courses for GO SLIM Biological Processes were constructed for the oligodendrocyte lineage using the 5 most significant terms. Pathway time course was represented applying the same method and using selected key pathways from the prior analysis.

Due to the potential reversibility of early (i.e. NSC) cell state transitions (Basak et al., 2018; Calzolari et al., 2015; Obernier et al., 2018), a considerable overlap among cell clusters in terms of expression profiles, Reactome and PANTHER Pathways, and Protein Class led to the definition of 3 broader groups. For this, the earlier stages constituting OLqNSCI, OLqNSCII and OLpNSC, the mid stages represented by OLaNSC and OLTAP, and the remaining OPCs and mature oligodendrocytes as the later stages were assembled. Results for signaling Pathways, Reactome and Protein Classes enrichment corresponding to the 3 groups were merged and the top third most significant terms for each of the 3 groups were charted as dot plots and ranked in descending order based on their enrichment. Results for transcription factor Protein Class for the 7 oligodendrocyte lineage clusters were visualized with a heatmap representing the top 3 most enriched PANTHER Pathways related to “transcription factor function” across the 7 clusters. The heatmap was obtained with the pheatmap package in R.

### Characterizing transcription factors for gene regulatory network (GRN) reconstruction

Once the most significant set of genes is determined, it is of importance to investigate if and how the genes are interacting at the molecular level. In addition to the genes previously identified as transcription factors, additional transcription factors known to play a central role in NSC differentiation were selected and their putative targets taken from http://genome.gsc.riken.jp/TFdb/tf_list.html, http://www.tfcheckpoint.org/index.php/browse, and PANTHER. Given the large numbers of transcription factors identified, a prioritization procedure was developed, relying on functional and transcription factor-transcription factor interaction data. Initially, the Cytoscape app GeneMANIA was used to identify functional interactions among the input genes (Franz et al., 2018). The data available for GeneMANIA for human gene orthologs are significantly more numerous and better characterized, therefore, mouse gene symbols were converted to human gene symbols. The genes used as input for GeneMANIA were limited to transcription factors and the queries were performed using the default parameter besides the type of interactions. In fact, the selected databases were “genetic” for downstream gene regulation and “physical” for protein-protein interactions. A first network (precursor to the one depicted in Figure S7) was thus obtained, on which additional analyses were performed for prioritizing the large numbers of transcription factors included. For this, the heat diffusion algorithm was applied (Carlin, Demchak, Pratt, Sage, & Ideker, 2017) to facilitate identification of central transcription factors according to their level of interactions: the higher the number of interactions, the higher the calculated heat diffusion rank. The heat diffusion algorithm was applied 3 times separately, once focusing on genetic interactions, (Figure S7b), once on physical interactions (Figure S7c), and finally combining the newly obtained heat diffusion ranks (Figure S7a). The diffusion ranking values so obtained were plotted against the number of interactions of each transcription factor. As additional information, the nodes size of the included transcription factors was adjusted to their combined physical and genetic interaction ranks in Figure S7d.

Aiming to group the transcription factors to organize the network similarly to the clustering of FateID, the precursor network was organized to depict transcription factor expression according to their stage of expression. The primary goal is focusing on oligodendrogenesis, hence transcription factors whose annotation showed an enrichment towards neuronal lineage only were discarded. Genes common to both lineages at different stages were processed in order to define transcription factors that show specificity to either lineage. The specificity threshold was defined at linear 2-fold. This selection enabled emphasizing genes/transcription factors that are expressed in the oligodendrocyte lineage still including those ones sharing neuronal signatures. To assign transcription factors to the appropriate oligodendrocyte or neuronal clusters, the principle of higher expression was adapted. Nodes and labels sizes were organized based on the FDR values. A separate grouping of transcription factors was made by selecting the transcription factors overlapping between aNSCs and TAPs. Transcription factors that were significantly expressed in at least 2 different stages of the oligodendrocyte lineage were classified as “Pan transcription factors”. GeneMania networks presented in Figure S7a were used as visual aids for facilitating the positioning of transcription factors according to the stages in differentiation they are expressed in.

### TETRAMER reconstruction of GRNs of SVZ lineages

The above step focuses on transcription factor functions in terms of their genetic transcription factor-transcription factor interactions. This information was used as the framework for a comprehensive inspection of transcription factor-target gene interactions. For unravelling GRNs, TETRAMER (TEmporal TRAnscription regulation ModellER) reconstructs fate transition-specific GRNs by integrating transcriptome data (user provided) with inbuilt information (Cholley et al., 2018) (available gene expression profiles, human genome-wide promoters and enhancers maps, and ChIP-Seq available datasets) (see also http://ngs-qc.org/tetramer/.). The input data for TETRAMER was constituted of a matrix containing 1467 genes – including transcription factors – irregularly expressed in the oligodendrocyte lineage during differentiation and, at the same time, expressed more in the neuronal lineage. Part of the genes were included in the input matrix due to their association with GO Biological Processes identified as major hallmarks of later stages of oligodendroglial maturation. These hallmark GO Biological processes were: (i) Signal Transduction, (ii) Ca Ion Transport, (iii) Synaptic Organization, (iv) CNS Development, and (v) Multicellular Organismal Development. All of them presented a *p*-value < 0.0002 (as estimated with Hypergeometric test). Specific hallmarks for mature oligodendrocytes in the considered GO Biological Process were: (i) Cytoskeleton Organization, (ii) Myelination, (iii) Myelin Maintenance, (iv) Cell Adhesion, and (v) Transport. All of them presented a *p*-value < 0.007 (as estimated with Hypergeometric test). Genes consistently enriched more than 2.5 folds (aside from transcription factors) in OPCs and mature oligodendrocytes in the lists above and among DEGs found with Seuratwere considered for a further functional enrichment analysis using the freely available Functional Enrichment analysis tool (FunRich) http://funrich.org/index.html. GO Biological terms were reduced to fit, and a complete list has otherwise been made publicly available (see below).

The following settings were necessary prior to assessing the transcriptional signaling propagation in the GRNs with TETRAMER. The matrix consisting of 1467 genes, among which are transcription factors expressed in both linages (common transcription factors) or exclusively in either lineage, was used to construct 3 separate subnetworks (presented in Figure 4b-d). These networks were filtered accordingly to highlight transcription factors expressed in the neuronal lineage and excluding genes expressed in OPCs and mature oligodendrocytes. The nodes expressed in the qNSCI stages were named “start node” as they represent the initial time point of differentiation, and transcription factors expressed in more than 2 stages were termed “pan transcription factors”. The nodes expressed in final stages of differentiation, i.e. OPCs and mature oligodendrocytes for the oligodendrocyte lineage, were named “end node”. The time course for differentiation was arranged in order from qNSCI to mature oligodendrocytes, and all intermediate differentiation stages were selected for network propagation. In the TETRAMER results section, all transcription factors containing a yield were considered to be part of the main signaling network. When the signal propagation network viewing was initiated, nodes in the network were colored from white/light red for the qNSCI nodes to darker red for mature oligodendrocytes, white to dark grey for transcription factors expressed in both lineage as ”common transcription factors”, and white to dark blue for transcription factors expressed in the neuronal lineage. Node sizes in these networks and subsequent analysis were graded based on the heat diffusion rank combined with additional parameters: node stress (calculated as the number of shortest paths passing through the considered node) indicating the extent of node activity, neighborhood connectivity, and the numbers of transcription factor-target gene interactions (edges). The values for each of these criteria were normalized in the range (0, 10) (low to high attribute values) and an average of all parameters was computed generating a final score termed as the connectivity index.

**Figure 4.**
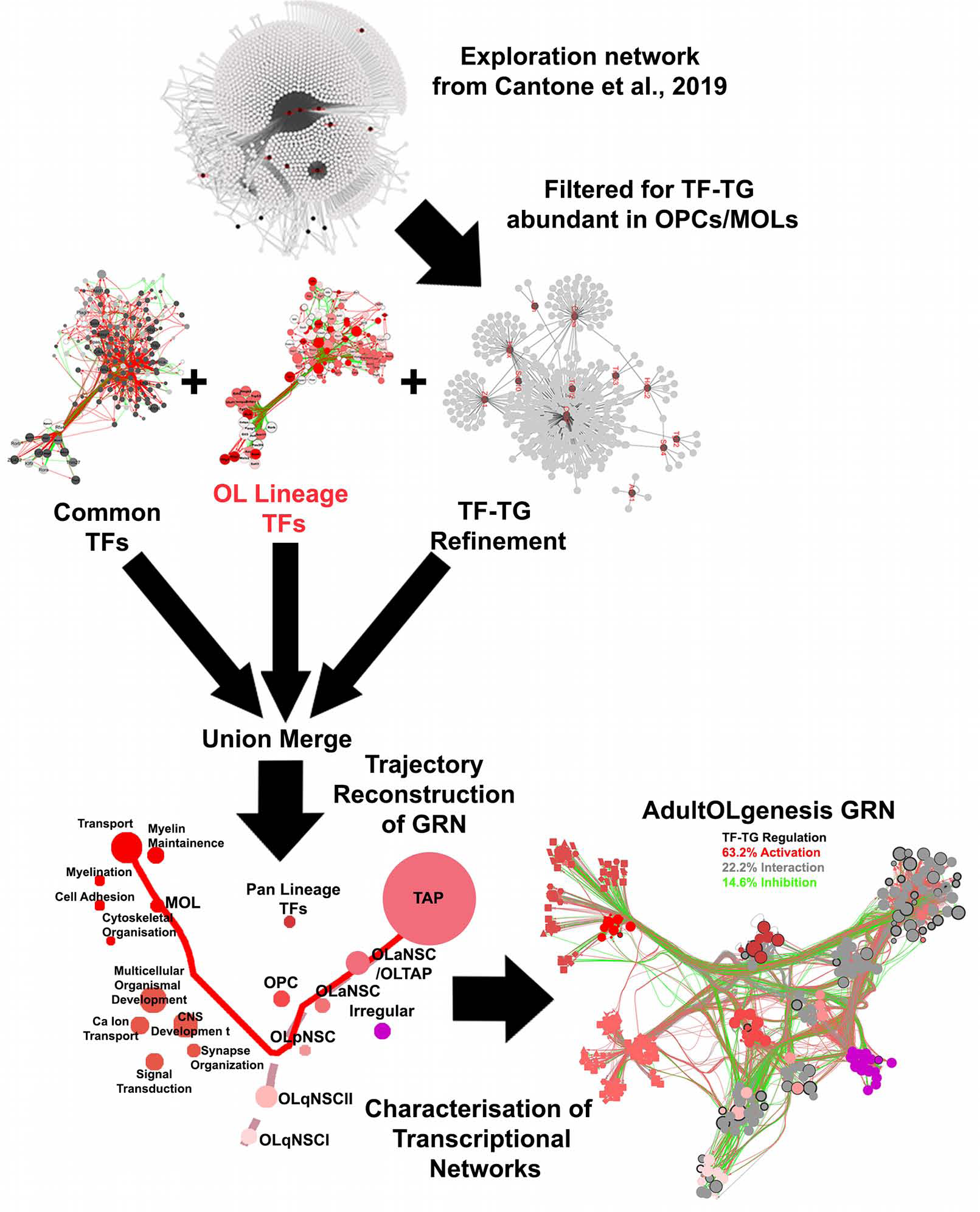
Reconstruction of a GRN by merging of previously generated networks and organization of stages in the oligodendrocyte lineage according to their pseudotime coordinates. Networks assembled in Figure S6b and d, those published recently (see description in results section) and additional transcription factor-target gene (TG) interactions from CHIP-seq studies are processed for a larger network. transcription factor expressed in 7 distinct stages are arranged according to their pseudotime order and each node represents the relative numbers of transcription factors expressed in a given phase in the lineage. A separate grouping of transcription factors expressed in both aNSCs and TAPs are plotted according to the average trajectory coordinates between these 2 phases. The top 5 GO Biological Processes in OPCs and mature oligodendrocytes are incorporated for identifying their regulatory pathways. Irregularly expressed transcription factors are those which are abundant in the neuronal lineage and additionally expressed along the oligodendrocyte lineage. The entire GRN of the oligodendrocyte lineage, termed as “AdultOLgenesis GRN”. Nodes in bold outer surface are characterized by high protein-protein interactions. Green, red and grey interactions signify gene inhibition, activation and unspecified interaction onto their target genes (TGs), respectively. Legends for the AdultOLgenesis GRN are presented in Figure 5 and 6. Abbreviations: aNSC: activated neural stem cell; GRN: gene regulatory network; MOL: mature oligodendrocyte; NB: neuroblast; NSC: neural stem cell; OLaNSC: oligodendroglial activated neural stem cell; OLpNSC: oligodendroglial primed neural stem cell; OLqNSCI: oligodendroglial quiescent neural stem cell (subtype I); OLqNSCI: oligodendroglial quiescent neural stem cell (subtype II); OLTAP: oligodendroglial transiently amplifying progenitor; OPC: oligodendrocyte precursor cell; TF: transcription factor; TG: target gene.

### Reconstruction of the AdultOLgenesis GRN

The GRNs representing transcription factors expressed in the oligodendrocyte lineage and the ones common to both SVZ lineages were examined further by combining data of these two networks. The input matrix used for TETRAMER was expanded with genes that account for the GO Biological Processes for later stages of oligodendrogenesis. The TETRAMER analysis was performed as described above. The newly obtained larger network was expanded further to include the interactions with transcription factors known to play a central role in NSC differentiation. Additional ChIP-seq data for *Ascl1, Cebpa, Hdac2, Jun, Sox4, Sox9, Tgif2*, and *Tpr53* were obtained from the Chip Atlas database (Oki et al., 2018), while *Hopx* (He et al., 2016; Jain et al., 2015), *Zeb1* (Rosmaninho et al., 2018) were taken from individual studies. For *Trp53* and *Zeb1* original data, the potential targets were annotated using human species gene names, hence to merge the results of the different datasets coherently, an R script was compiled for converting human-annotated to mouse-annotated genes (biomaRt package) (Durinck, Spellman, Birney, & Huber, 2009). For the potential gene targets extracted from the Chip Atlas database, a threshold on the score returned by the database was defined to select the targets with highest level of confidence (threshold values: 250). For *Hopx*, a different thresholding approach was applied, for the available data were processed differently: ChIP-seq data were processed with the Homer suit of tools, and therefore a different score was assigned to each identified peak. To discriminate between the relatively reliable targets, the average score was calculated and set as the discriminant threshold. For all the added transcription factors, a node was added to the precursor GRN representing the transcription factor itself. Interactions representing the ChIP-Seq data were created between the transcription factor and the targets already included in the precursor GRN. In order to select the target genes and generate this representation, a Python script was compiled. The code generated a text file associating to each transcription factor-target pair the source of the datum, the type of the interaction, and the type of the source data. To extend the number of interactions of the GRN specifically or oligodendrogenesis, the transcription factor-target gene networks were imported independently in Cytoscape and union-merged with the Exploration network recently published by Cantone and colleagues (Cantone et al., 2019). The union merge was repeated between the latest union merge and the AdultOLgenesis Network generated by TETRAMER, thus incorporating additional information. For the 11 selected transcription factors, targets were selected based on (i) a score describing the confidence of the actual formation of the transcription factor-target gene binding, thus increasing the reliability of the interaction, and (ii) the biological context of the transcription factor-target gene binding by limiting the set of genes to the ones previously identified in the Core GRN. The confidence in the identification of the same transcription factor-target gene binding in human and mouse species is ensured by a high level of genomic similarity.

The obtained network was remodeled for positioning the nodes according to their pseudotime trajectory coordinates, thus providing an improved overview for transcriptional processes. The GeneMania network presented in Figure S7a was used as reference to guide the correct placement of genes in individual stages of differentiation. The Cytoscape plugin CoordinatesLayout version 3.0 and copycatLayout version 0.0.9 were used. The average coordinates for defined cells at each stage of differentiation was calculated along its x-y axis, and at each of the estimated positions, a node was added to represent the corresponding differentiation stage. The size of the node was scaled to reflect the relative abundance of transcription factors associated to the differentiation stage. The “mock” nodes representing oligodendrocyte cell clusters were organized according to their FateID trajectories. Nodes representing transcription factors/genes associated with each of the different stages of the oligodendrocyte lineage were grouped together and force directed using Compound Spring Embedder (CoSE), and overlaid with the mock nodes. Settings for CoSE were adjusted systematically for all oligodendrocyte stages with edge lengths kept constant and spring strength/repulsion adjusted equally. Nodes with higher heat diffusion ranks were permitted to overlay those genes with lower heat diffusion ranks to highlight transcription factors with greater functional relevance. Interactions were bundled using handles with the following parameters: 0.003 spring constant, compatibility threshold of 0.3 and 500 maximum iterations. Nodes without interactions (that is, without edges) were excluded and hidden in the background. Summaries highlighting downstream transcriptional regulation (outwards transcriptional propagation) were constructed by adjusting node and edge sizes reflecting the relative numbers of transcription factors/genes in the different stages of differentiation and the downstream activation or repressive transcriptional control, respectively. Five additional GO Biological Processes were manually included close to later-stage oligodendrocytes. The entire reconstructed network was termed “AdultOLgenesis GRN” and was thus an expansion of a recent study describing GRNs during OPC-to-mature oligodendrocyte differentiation (Cantone et al., 2019) by inclusion of additional regulatory features.

### Assessing transcriptional propagation by selected transcription factors and Signaling Pathways

Transcription factors expressed in defined stages of oligodendrogenesis (see also Figure S7) or associated in PANTHER pathways were tested for transcriptional propagation across the Core GRN (see also Section 7 of the online repository: https://github.com/kasumaz/AdultOLgenesis). The transcription factors were selected with their direct target genes and subsequent secondary and tertiary target genes. All primary target genes were retained, whereas secondary and tertiary target genes, if they were transcription factors, were filtered according to their protein-protein interaction and connectivity index of >7.5 and >7, respectively. Biological Process GO terms were preserved throughout. Interactions of transcription factors associated with PANTHER pathways, secondary and tertiary target genes were sized in order of line thickness in this order. Manual curation was operated for correcting the edges between GO Biological Processes and PANTHER pathways. Transcription factors in each network were ranked in heatmaps for each pathway, according to the number of target gene interactions (activation/inhibition) across the different pathways selected for analysis. Gene regulatory data for the quantification of activation, inhibition and unspecified interaction were obtained from the edge section of the AdultOLgenesis GRN in Cytoscape.

## RESULTS

### Identification of the earliest stages of the SVZ-derived oligodendrocyte lineage

Transcriptional networks controlling oligodendrogenesis from adult SVZ-NSCs are currently unsolved. Here, NSCs/progenitors and subtypes of neural cells found within the young adult SVZ were isolated and FACS-enriched using genetic reporters and immunochemical markers, as described in a previous report on adult neurogenesis (Basak et al., 2018) (Figure S1a). Relying on well-established markers for the earlier stages in oligodendrogenesis as a guide, we identified putative lineage-specific subpopulations of NSCs (Figure S1b). Approximately 1200 single cells were visualized using tSNE plots and used for categorizing previously defined clusters and subclusters (Basak et al., 2018). We hypothesized that, as in the early postnatal SVZ, adult lineage-specific NSCs could be identified by their expression of oligodendroglial and neuronal hallmark genes. To this end, previously generated lists of genes associated with SVZ-oligodendrogenesis from bulk datasets were assembled and searched for oligodendrocyte lineage-enriched genes when compared to neurogenic populations (Azim et al., 2017). In this initial step, PartekFlow was used to input a number of landmark genes that define the earlier stages of the oligodendrocyte lineage. These include the essential transcription factors *Olig2*, *Olig1* and *Sox10*, and a cohort of other transcription factors as described in Materials and methods (Figure S1b). As an advantage, PartekFlow highlights cells based on input gene marker profiles, allowing the identification of neighboring cells that are transcriptionally similar. Following classification of putative oligodendrocyte lineage cells descending from NSCs/TAPs, gene lists derived from PartekFlow had initially revealed that the major differences in the 2 SVZ lineages are attributed to the expression of transcription factors (see below), in agreement with the observed pre-eminence of transcription factors as lineage-specific markers within postnatal gliogenic lineages (Azim et al., 2015). Thus, expression levels of transcription factors were explored. In particular, single NSCs/TAPs expressing the above 3 essential transcription factors, as well as a cohort of others (Figure S1b) expressed in postnatal gliogenic NSCs were identified as likely to contribute to the oligodendrocyte lineage (Figure 1a) and amounted to approximately 7.2% of all NSC/TAPs across the 5 NSC and TAP stages of differentiation, which is in line with previous reports describing the relative scale of SVZ-oligodendrogenesis when compared to the generation of neurons (Menn et al., 2006). This particular subpopulation is identified by the co-expression of the known pan-NSC marker *Hes5* in qNSCs I/II expressing *Olig2* (Figure 1b). The first 3 stages of identified OLNSCs also expressed the pallial marker *Pax6* whereas they did not express the ventral SVZ markers *Gsx2* and *Nkx2-1* (Figure 1b). Most notably, an initial analysis of differentially expressed genes (DEGs) revealed patterns that are reminiscent of transcriptional programs found during postnatal oligodendrogenesis (Figure S1b). Identified cells of the oligodendrocyte lineage in the adult cluster closely with corresponding pooled populations (qNSCI-pNSC) in transcription factor expression, including postnatal gliogenic NSCs/TAPs (Figure S1c), reinforcing the view that subsets of adult NSCs are committed to an oligodendrogenic fate.

Next, FateID pseudotime analysis was performed to assess whether identified putative oligodendrogenic and neurogenic NSCs define distinct cell state trajectories (Herman et al., 2018). The genes that are significantly varying as determined by the Seurat analysis (see Materials and methods) in oligodendrogenic and neurogenic lineages, and the genes enriched in mature oligodendrocytes, were used for testing the hypothesis that the selected oligodendrocyte lineage cells engage in a trajectory from earlier subpopulations of NSCs/TAPs. Oligodendrocytes and neuroblasts were used as the final lineage endpoints. Neuroblasts followed a continuous trajectory from qNSCI and many neurogenic TAPs resembled neuroblasts in pseudotime (Figure 1c). Addition of mature oligodendrocytes to this mostly neurogenic trajectory forces an early branching for the oligodendrocyte lineage, at qNSCI/II stages. However, when considering mature oligodendrocytes alongside putatively oligodendrogenic NSCs/TAPs and OPCs, mature oligodendrocytes branched off the main trajectory closer to OLaNSCs, TAPs and OPCs. Interestingly, addition of neuroblasts to the oligodendroglial trajectory resulted in a transcriptional path that overlapped partly with fewer OLTAPs compared to the neurogenic TAPs, highlighting the distinction between the two lineages. Genes detected in early-to mid-stage oligodendrocyte lineage cells (qNSCI to TAPs; Figure 1d-f) regulate signaling-to-transcriptional control of the oligodendrocyte lineage. Cluster-enriched markers (Figure 1f) include genes coding for the G-protein-coupled receptor genes *Lrig1* and *Adora2b* for OLqNSCI (Poth, Brodsky, Ehrentraut, Grenz, & Eltzschig, 2013; Simion, Cedano-Prieto, & Sweeney, 2014); the Bmp4-responsive gene *Htra1* serine protease (Chen et al., 2018), and the gliogenic transcriptional adaptor *Hopx* for OLqNSCII (Zweifel et al., 2018); the transcriptional regulator *Foxo1* (Kim, Hwang, Muller, & Paik, 2015), or the mitochondrial ketogenic enzyme *Hmgcs2* in OLpNSCs (Jebb & Hiller, 2018). OLaNSC and OLTAP markers include genes encoding for *Hspa1a* and *Hspa1b*, and *Cdk4* and *Hes6*, respectively (Arion, Unger, Lewis, Levitt, & Mirnics, 2007; Kim et al., 2015; Lukaszewicz & Anderson, 2011). This analysis revealed that many of the cluster-specific genes belong to families involved in signaling and transcriptional control, and demonstrated that cells representing the earliest stages of the oligodendrocyte lineage are identifiable among NSCs, allowing the analysis of lineage-specific signatures. Further examples of pan lineage markers are given in Figure S2.

### Stage-specific processes

Following determination of stage-/cluster-specific signatures, we focused on individual lineages (oligodendrocyte and neuronal, separately) and analyzed the cell clusters in each (7 oligodendroglial clusters, 6 neuronal clusters) for pathway enrichment using the PANTHER database. The most significant pathways for each stage of the 2 lineages are illustrated in dot plots showing the activity of a given pathway across the different clusters (Figure 2a). In this analysis, the genes exclusively belonging to 1 cluster were examined. Interestingly, the earliest stages of the oligodendrocyte lineage (qNSCI-pNSC) were uniquely characterized by the activity of pathways (e.g., “angiogenesis”, “cysteine biosynthesis”, “cytoskeletal regulation by RhoGTPases” and “pyruvate metabolism”), which also distinguished them from early NSC stages of the neuronal lineage. Contrastingly, a considerable overlap in pathways in the mid-stages (aNSC and TAPs) between the 2 lineages was evident, with few exceptions such as “Notch signaling” that was prominently enriched in OLaNSCs, and “p53 and cell cycle pathways” enriched in the same stages of the neuronal lineage.

All genes enriched in the oligodendrocyte lineage clusters (including those shared with the neuronal lineage) were next examined for characteristic biological processes (Slim Biological Processes in PANTHER) and plotted across the various stages of lineage progression. Machineries related to “lipid or energy metabolism”, “response to signaling” or “developmental patterning molecules” were enriched in the earlier stages, whereas mechanisms such as “translation”, “gene expression” and “cell division” were most prominent in OLaNSCs and OLTAPs (Figure 2b), consistent with previous observations (Llorens-Bobadilla et al., 2015). At a later stage of differentiation, OPCs engage in processes that were also active in OLqNSCII. For example, GO biological process “cell communication” is enriched in OLqNSCII, and then again at the OPC stage (Figure 2b), possibly accounting for the close proximity of these 2 segregated populations in both the tSNE plots and pseudotime (Figure 1a,c). Many of the characteristic processes detected in mature oligodendrocytes increase in expression during the course of lineage progression. We expanded this analysis for both protein class and signaling pathways, to include the neuronal lineage (Figure S3), observing that transcription factors and nucleic acid-binding proteins are highly abundant in the early and mid-stages of the oligodendrocyte lineage along with major developmental pathways. A time course for the expression of the most significant signaling pathways in this analysis points to key processes regulating transition along the oligodendrocyte lineage (Figure S4), with increasing transcriptional activity suggested by the pattern of expression of transcription factors and nucleic acid-binding proteins across the different oligodendrocyte lineage stages (Figure S5). Altogether, these findings outline potential mechanistic differences in the control of progression within the 2 SVZ-derived lineages, with notable differences in inferred signaling pathway activities and in the expression of transcriptional modulators.

### Cell cycle properties of SVZ-NSCs/progenitors

Given the observed differences in cell cycle, transcriptional control and nucleic acid synthesis pathways in mid-stages (OLaNSC, OLTAP) of the oligodendrocyte lineage we further explored stage-specificity. To this end, cell clusters were analyzed by computing cell cycle phase scores based on canonical markers for the 3 main cycle phases (S, G2M and G1 phase) using the Seurat V3 package. Single cellś expression levels of established cell cycle markers were compared and data were regressed out by modeling the relationship between expression signatures of each cell with cell cycle markers. Upon identification of the predicted cell cycle state for cells of all 13 clusters, those representing the oligodendrocyte lineage were plotted in accordance to their pseudotemporal coordinates and summarized in bar plots for both lineages (Figure 3a,b), and signal intensities as ridge plots (Figure 3c). In both lineages, the early stage NSCs (qNSCI-pNSC) and OPCs were almost entirely at the G1 phase with signal intensities shifted towards baseline, whereas S and G2M marker signals were detected in neuroblasts (Figure 3c). More than half of the aNSCs in either lineage were actively cycling and rarely at the G2M phase, whilst the remaining were quiescent. Although no proneuronal TAP was found to be quiescent, with just over 55% at the G2M phase and the remaining in S phase, a slightly greater proportion of OLTAPs were in S phase. Signal intensities for the S and G2M markers were relatively absent in the early neuronal lineage.

Following identification of the mid-stage cell cycle properties, Seurat V3 was used to identify cluster-specific genes in OLaNSCs and OLTAP by subclassification according to their predicted cell cycle phase. Therein, transcripts significantly expressed by the 9 cell clusters (OLqNSCI, OLqNSCII, OLpNSC, OLaNSC in G1, OLaNSC in S Phase, OLTAP in G2M, OLTAP in S Phase, OPC and mature oligodendrocytes) were extracted and a further pathway analysis was performed on the mid-stages of the oligodendroglial lineage (OLaNSC in S Phase, OLTAP in G2M and OLTAP in S Phase). Pathways and Protein Class determination was performed for OLaNSCs in the G1 and S phase in the intercept (genes overlapping in the 2 groups) in this analysis and the same for G2M and S phase for OLTAPs (Figure 3d, e). Interestingly, aside from Notch and Hedgehog signaling, metabolic-like mechanisms that include thiamine metabolism and threonine biosynthesis are also features of quiescent OLaNSCs, whereas cycling aNSCs were characterized by pathways regulating transcription, translation, and nucleic acid binding. In OLTAPs at the G2M phase, EGF-, p38 MAPK-, p53 and interferon gamma-signaling were significantly enriched. Furthermore, as expected, many of the pathways expressed in actively cycling TAPs included regulation of mRNA, replication, and cell cycle-related processes (Figure 3d). Cell cycle-related analysis of mid-stage oligodendrocytes accordingly revealed that the major protein classes expressed in G2M and S phases were “nucleic acid-binding” and “transcription factor”, whereas OLaNSCs in G1 were characterized by extracellular matrix and vesicle-associated SNARE proteins (Figure 3e). Altogether, examining heterogeneity of cell cycle states in proliferative cells of the oligodendroglial lineage suggests that transcriptional cues are a major driver in OLqNSCII and in mid-stages of the lineage, which warrants further investigation.

### Gene regulatory networks in NSCs/TAPs driving myelination

The previous analysis disclosed major classes of genes expressed in earlier oligodendroglial lineage cells, notably with mid-proliferative cell stages featuring abundant expression of transcriptional modulators. Therefore, the expression of transcription factors (including transcriptional adapters and transcription cofactors) was further explored in both the oligodendroglial and neuronal lineages. Transcription factors were prioritized via a GeneMania-based genetic and physical interaction analysis. This step facilitates predicting the effects of genetically perturbing a given transcription factor, in terms of its impact on neighboring genes, based on predicted transcription factor-to-targetgene interactions and protein-protein interactions (see Materials and methods; Figure S6). To determine which transcription factors are most relevant according to their genetic perturbation, genes were prioritized by applying the “heat diffusion” algorithm that tests the input query data and the functional interaction of each gene for propagation across the network. Strongly connected nodes in the network, supporting sufficient regulatory activity propagation, are uncovered and allow creation of subnetworks, enabling exclusion of nodes with lower functional relevance. Prioritization steps were applied for both physical (protein-protein interactions) and genetic interaction (Figure S6a and expanded further in Figure S7) and identified transcription factors with potentially central roles in specifying cell states.

We next employed TETRAMER, a tool integrating inferred differentiation states during lineage progression with gene regulatory networks (GRNs) from publicly available datasets (Cholley et al., 2018), for the reconstruction of GRNs whose activity may define cell states within the oligodendroglial and neuronal lineages. Briefly, lists of transcription factors expressed in the identified clusters of the oligodendroglial and neuronal lineages as well as transcription factors that are expressed in both of these lineages (“common” transcription factors), were used as input by TETRAMER to explore gene expression profiles of mammalian tissues/cells with comprehensive analysis of deposited ChIP-sequencing data corresponding to transcription factor-binding and further comprising data on actively transcribed enhancers and promoters. In the preliminary step, transcription factor-transcription factor and transcription factor-target gene interactions were compiled for determining regulatory interactions between shared neurogenesis- and oligodendrogenesis-restricted (Figure S6b-c) transcription factors. A large proportion (89.3-91.8% of all interactions) of transcription factor-target gene interactions for transcription factors common to both lineages and those regulating neurogenesis consisted in predicted target gene activation. Interestingly, however, activating and inhibiting regulatory interactions were predicted to occur at similar frequencies between oligodendrogenesis-specific transcription factors, highlighting qualitative differences in how cell state transitions are controlled at the transcriptional level in the 2 lineages. Transcription factor interactions with target genes were quantified using the Cytoscape network analysis parameters by probing the entire training GRN reconstructed (as described in Figure 4) and presented as intensity heatmaps for the top 25 transcription factors with the most global, activating and inhibiting target gene interactions in Figure S6e (ranked transcription factor examples in Figure S6f). Amongst the most highly ranked transcription factors were *Cebpa,* expressed in OLqNSCII and best described in the context of myelopoiesis, where it inhibits proliferation (see (Calella et al., 2007) for proposed roles during neural development), and *Ezh2*, which promotes embryonic NSC proliferation and differentiation into OPCs (Sher et al., 2008). Altogether, these analyses enable ranking transcription factors that are likely to act as “master regulators” based on a number of parameters that include their target gene regulation, protein-protein interactions and diffusive effects downstream. To further investigate the major gene regulatory cascades controlling adult oligodendrogenesis, the GRN assembled for common transcription factors and oligodendrogenesis (Figure S6b,d) were combined and merged with a recently assembled GRN for post-OPC stages of oligodendroglial maturation (Cantone et al., 2019) (Figure 5), which defined the role of transcription factors such as *Olig2*, *Sox10*, and *Tcf7l2* during oligodendrocyte differentiation and maturation. Here, genes overlapping in later-stage oligodendrocytes from the present study with those published recently (Cantone et al., 2019), were filtered and represented as GO Biological Processes in a larger merged network (Figure 5). Approximately, a quarter of the differentially expressed genes, as determined by the Seurat analysis, were used in this TETRAMER analysis. Of note, modulation of many of these genes has previously been reported during postnatal and adult oligodendrogenesis from NSCs (Azim et al., 2015). This final network termed as “AdultOLgenesis GRN” was used for further analysis as described below. The individual stages were then positioned relative to the average positioning of single cells/clusters in FateID pseudotime (Figure 1c) (see Materials and methods for network assembly).

**Figure 5.**
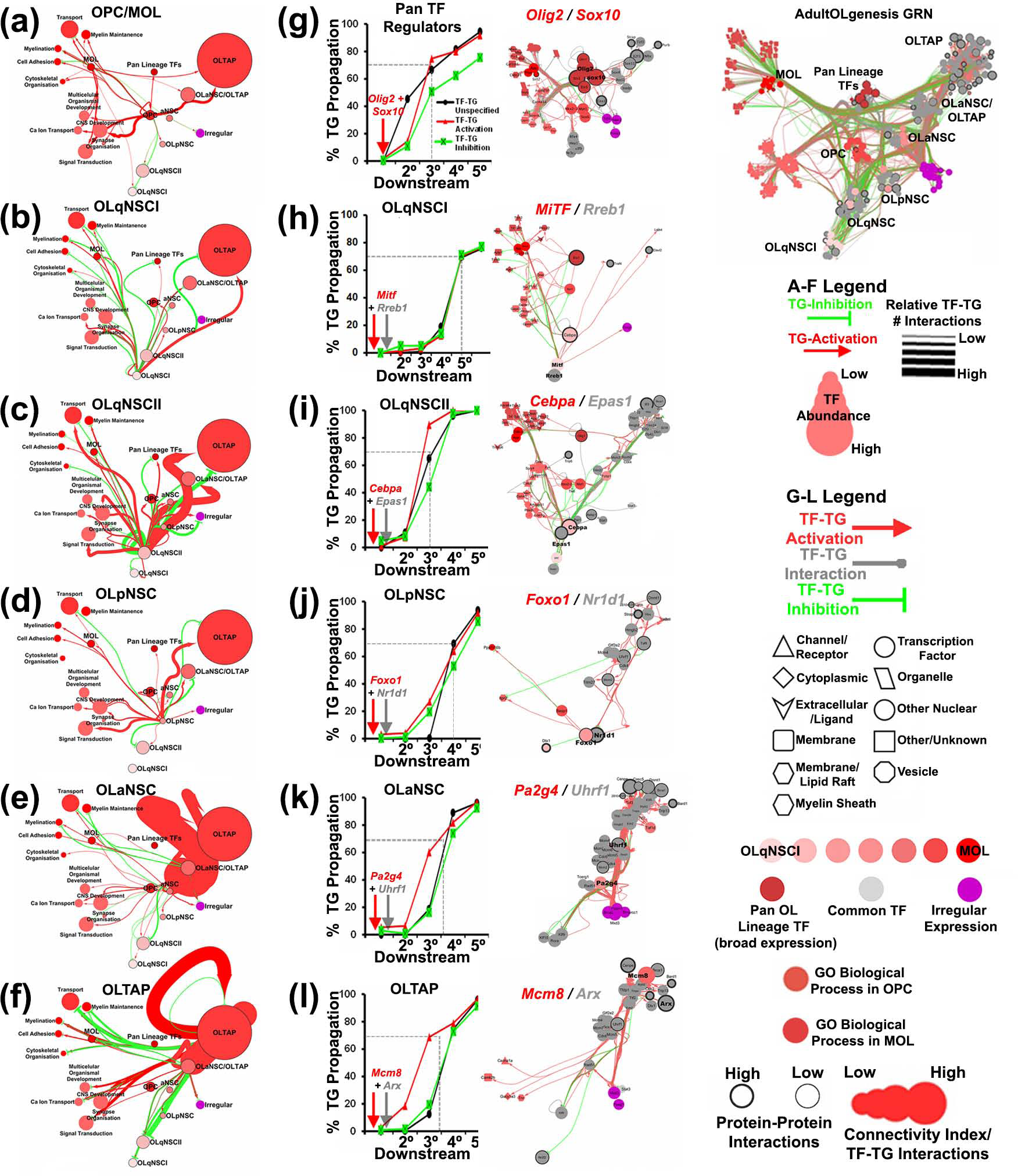
Gene regulatory interactions in substages of the oligodendrocyte lineage and the predicted effects of key transcription factors in each stage. (a-f) Summaries of the GRNs in each stage showing downstream gene regulation propagation. Edges and nodes reflect the relative numbers of gene regulatory interactions and numbers of transcription factor expressed in each stage, respectively. (g-l) The downstream gene regulatory interactions of selected transcription factors are quantified by propagating the transcription factor most highly ranked in the oligodendrocyte lineage and a transcription factor that is commonly expressed in a defined stage. In g, *Olig2* and *Sox10* are probed for their downstream target (designated as secondary, 2°), the subsequent downstream targets (including both genes in the GO Biological Processes and additional transcription factors) to the tertiary genes (3°). The transcription factor-target interactions quantified are plotted as a percentage versus the total numbers of genes/gene regulatory interactions from the entire reconstructed GRN. For each stage, the 2 highly ranked transcription factors and their direct target genes are shown as subnetworks from the entire GRN. Abbreviations: aNSC: activated neural stem cell; GRN: gene regulatory network; MOL: mature oligodendrocyte; NB: neuroblast; NSC: neural stem cell; OLaNSC: oligodendroglial activated neural stem cell; OLpNSC: oligodendroglial primed neural stem cell; OLqNSCI: oligodendroglial quiescent neural stem cell (subtype I); OLqNSCI: oligodendroglial quiescent neural stem cell (subtype II); OLTAP: oligodendroglial transiently amplifying progenitor; OPC: oligodendrocyte precursor cell; TF: transcription factor; TG: target gene.

### Stage-specific GRN

Focusing on target gene activation or repression in substages of the oligodendrocyte lineage reveals varying and contrasting degrees of gene regulation (Figure 5a-f). Notably, the OLqNSCII, OLaNSC and OLTAP stages exhibited prominent gene regulatory activities, with OLaNSCs seemingly progressively activating TAP-specific target genes (Figure 5b,e,f), whilst also inhibiting the expression of genes typical for the most differentiated (OPC/mature oligodendrocyte) stages. This suggests that, following activation, oligodendrocyte lineage cells would steadily progress to the TAP stage, which could be temporarily stabilized by active repression of differentiation-associated gene modules. Interestingly, the uniquely low expression levels of transcription factors active in OLqNSCI and OLpNSCs are predicted to only modestly result in GRN state restructuring at these stages. These findings demonstrate that transcription factors expressed in distinct stages during oligodendrogenesis have contrasting modes of gene regulation.

Next, the highest ranked transcription factors for each of the selected oligodendrocyte lineage stages were probed for the extent to which their inferred transcriptional regulatory activity propagates within the assembled GRN. The downstream gene regulatory effects of *Olig2* and *Sox10*, which modulate the expression of a large number of genes involved in gliogenic and neurogenic processes (Cantone et al., 2019), provided a quantitative reference to assess the GRN-modulating activity of newly identified transcription factors. The immediate downstream target genes for *Olig2* and *Sox10* are presented in a subnetwork (Figure 5g), and downstream propagation cumulatively modulates 75% of all network components within one further tier. The regulatory activity of *Mitf* and *Rreb1*, expressed in OLqNSCI, spread the least within the GRN, supporting the previous analysis (Figure S6e) that shows *Rreb1* as the highest ranked transcriptional repressor (Figure 5h). The impact of OLqNSCII transcription factors *Cebpa* and *Epas1* quantitatively resembles that of *Olig2* and *Sox10* (Figure 6i). Interestingly, *Cebpa* has been described for myelopoiesis as inhibiting cell cycle-related genes (Calella et al., 2007), which fits with its predicted inhibitory interactions with major cell cycle genes expressed in TAPs, supporting its role as a quiescent NSC-specific transcription factor. The 2 highest-ranked OLpNSC transcription factors, *Foxo1* and *Nr1d1*, induce the expression of a number of cell cycle regulators expressed in OLaNSCs and OLTAPs (Figure 5j). Like OLqNSCIIs, OLaNSCs and OLTAPs express transcription factors whose predicted influence on oligodendrocyte GRN activity resembles that exerted by *Olig2* and *Sox10*. These findings identify putative core regulators of stage-specific oligodendrocyte GRNs, thus complementing known pan-oligodendrocyte transcription factors *Olig2* and *Sox10* and providing a view into stage-specific control of oligodendroglial lineage progression.

**Figure 6.**
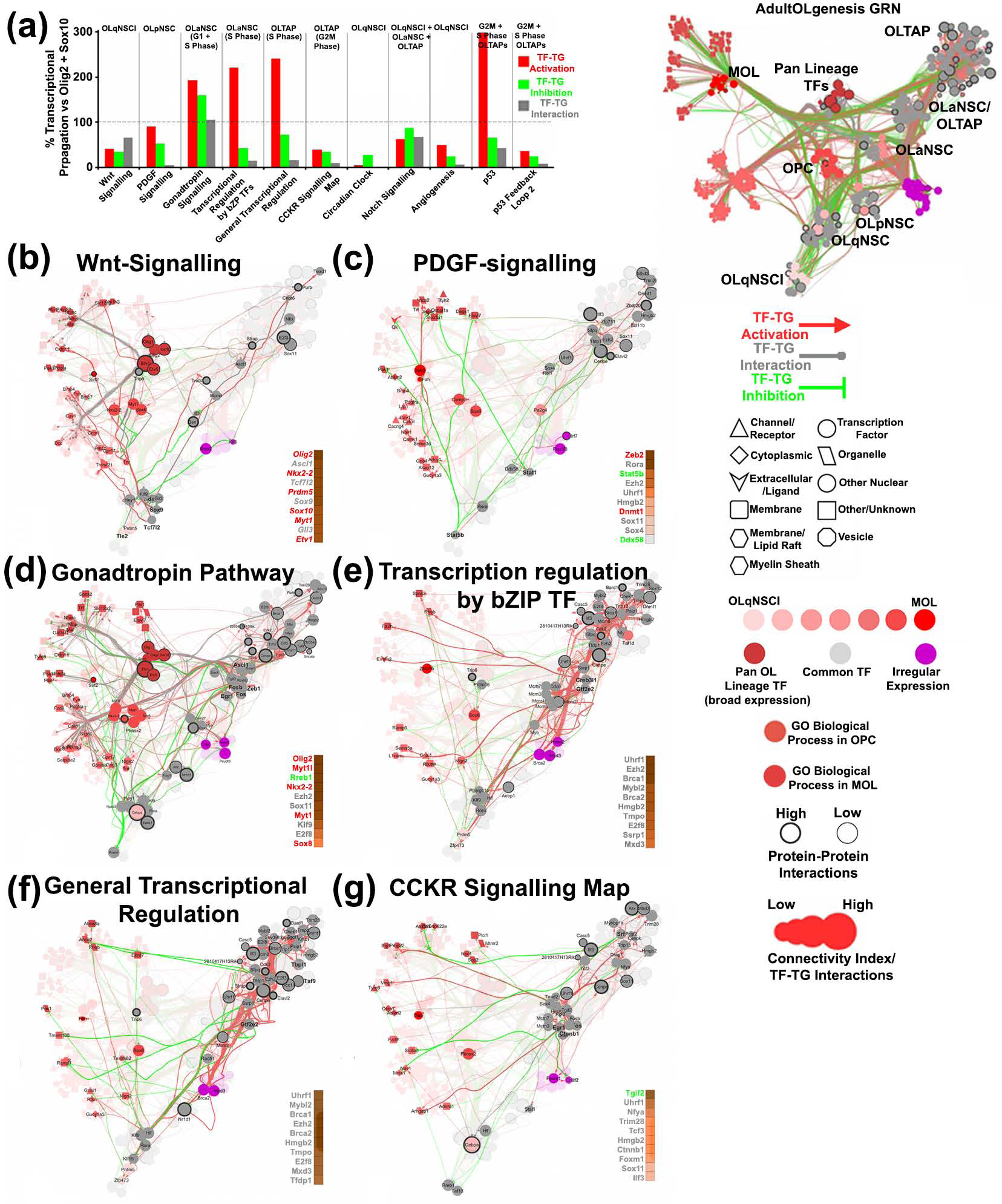
Mapping the impact of Signaling/PANTHER pathways onto GRNs. (a) PANTHER pathways from previous GO analyses were used to filter out transcription factors associated with each pathway term and their propagation onto the GRN were quantified and represented as a percentage compared to *Olig2*/*Sox10* (Figure 5g). (b-g) Examples of PANTHER Pathway effects onto the reconstructed GRN where dashed edges show the secondary effects of propagation compared to the direct effects of transcription factors associated with each pathway. Heatmaps show the relative ranking of key transcription factors of each pathway (both direct and secondary targets). Transcription factors highlighted in red, green and grey are those that activate expression of genes in later stage oligodendrocytes, inhibit the expression of genes in later stage oligodendrocytes and transcription factors that are essential to pathways as general regulators, respectively. Abbreviations: aNSC: activated neural stem cell; GRN: gene regulatory network; MOL: mature oligodendrocyte; NB: neuroblast; NSC: neural stem cell; OLaNSC: oligodendroglial activated neural stem cell; OLpNSC: oligodendroglial primed neural stem cell; OLqNSCI: oligodendroglial quiescent neural stem cell (subtype I); OLqNSCI: oligodendroglial quiescent neural stem cell (subtype II); OLTAP: oligodendroglial transiently amplifying progenitor; OPC: oligodendrocyte precursor cell; TF: transcription factor; TG: target gene.

### PANTHER Pathway Integration of GRNs

Having defined GRN state transitions controlling oligodendrocyte lineage progression, we attempted integrating this knowledge with stage-specific signaling pathway activity predictions from our previous PANTHER analysis (Figure 2b), reasoning that the identified signaling pathways would impact cell fate by ultimately modulating transcriptional networks. We explored this hypothesis using the reconstructed AdultOLgenesis GRN, examining signaling pathways known to modulate at least 2 transcription factors expressed at a given stage, and then assessing the predicted propagation of their transcriptional modulatory activity within the GRN. The direct target genes and additional secondary target genes regulated by both *Olig2* and *Sox10*, as performed in the prior analysis in Figure 5g, were used as a gauge for transcriptional coverage of each tested transcription factor associated with PANTHER pathway terms. In addition, the transcription factors induced by selected PANTHER signaling pathways were ranked based on the number of transcription factor-target gene interactions. These were further classified according to their gene-activating, -inhibitory, and unspecified effects onto the oligodendrocyte lineage in red, green, and grey, respectively (Figure 6a). In the earlier stages of the oligodendrocyte lineage, Wnt signaling is predicted to affect approximately half as many GRN components as those modulated by *Olig2* and *Sox10*. Our previous reports demonstrated that manipulating the Wnt signaling pathway by pharmacologic or genetic strategies increases the expression of *Ascl1*, *Olig2* and *Sox10* transcripts by at least 2.5-fold (Azim, Fischer, et al., 2014; Azim, Rivera, et al., 2014). In the present study, these essential transcription factors were indeed predicted to be secondary target genes of the Wnt pathway (Figure 6a,b), thus potentially accounting for the previously described enhancements of myelination following Wnt signaling modulation (Azim et al., 2017; Azim, Fischer, et al., 2014; Azim, Rivera, et al., 2014) and validating this strategy. PDGF signaling on the other hand induces similar gene regulatory events as *Olig2* and *Sox10*, although initially repressing genes expressed in later stage oligodendrocytes (Figure 6c), in line with its role as an OPC mitogen. However, *Stat1* and *Stat5b*, transcription factors modulated by the PDGF pathway, activate transcriptional networks in TAPs that enhance the expression of genes in OPCs and mature oligodendrocytes, leading to extensive downstream transcription factor induction that included a number of master regulators (see heatmap in Figure 6c). Interestingly, the pathways active in OLaNSCs (“Transcriptional regulation by bZIP transcription factors”, “Gonadotropin signaling pathway”) and in OLTAPs (“General Transcriptional Regulation”) are involved in a number of gene regulatory events larger even than those elicited by *Olig2* and *Sox10*, suggesting extensive GRN reorganization at these stages, possibly reflecting a lineage watershed on the way to differentiation. Indeed, a number of the above identified master regulators (Figure S6e) are key effectors of PANTHER pathway terms enriched in OLaNSCs and OLTAPs. Other potent signaling pathways examined include Notch signaling and p53 active in OLaNSCs (G1 and S phase) and TAPs (G2/M and S phase). Other pathways identified in the earlier PANTHER analysis, including “Angiogenesis” (OLqNSCI), “Circadian clock” (OLqNSCII/qNSCII) and “p53 Feedback Loops 2” (G2M and S phase OLaNSC/OLTAP), had overall fewer predicted transcriptional effects. Altogether, these findings demonstrate that large cohorts of transcription factors in the reconstructed Core GRN are effectors of multiple signaling pathways and provide useful insights into how environmental cues could affect transcriptional activities, and thus lineage progression, during SVZ-derived oligodendrogenesis. Future studies will aim at expanding these observations by testing the functionality of identified signaling pathways and to confirm changes in target expression experimentally.

## DISCUSSION

In the present study, a meta-analysis of single cell RNAseq data from the adult SVZ was performed to resolve the transcriptional signatures that distinguish between adult neurogenesis and oligodendrogenesis. Recent transcriptomic analyses focusing on early postnatal development and adulthood (Azim et al., 2017; Azim et al., 2015; Mizrak et al., 2019), provided some insights into the gene expression profiles associated with oligodendrogenesis. A number of newer transcriptomics studies of adult neurogenesis have shed light onto the molecular processes regulating the generation of olfactory bulb neurons (reviewed in (Marcy & Raineteau, 2019)). In addition, landmark studies using single cell profiling of murine oligodendrocytes from different stages of life in the CNS have immensely contributed to our understanding of the heterogeneity and molecular regulation of oligodendrocyte differentiation (Marques et al., 2018; Marques et al., 2016; Zeisel et al., 2015). A recent elegant single-cell sequencing survey of the lateral and medial walls of the young adult SVZ demonstrated that NSCs located in microdomains other than the lateral wall generate oligodendrocytes (Mizrak et al., 2019), but an in-depth investigation into the mechanisms controlling adult oligodendrogenesis had not yet been performed. To this end, we analyzed recently generated single-cell data from the SVZ (Basak et al., 2018) focusing on NSC/progenitor heterogeneity, in order to identify putative oligodendrocyte- and neuron-restricted progenitors (Felipe Ortega et al., 2013). Our findings reveal transcriptional regulators as major protein classes modulating all stages of the SVZ-NSC-derived oligodendrocyte lineage.

We carefully inspected the transcriptomes of NSCs and TAPs for the expression of known early oligodendroglial lineage marker genes (Figure S1). These included members of the Sox transcription factor family and *Olig2/Olig1*, which were found to be expressed selectively in NSC/TAPs that also expressed other known gliogenic markers such as *Hopx, Notch* genes and their subsequent target genes of the *Hes* family, overall resembling the pattern of expression observed during postnatal oligodendrogenesis (Azim et al., 2015). Oligodendrocyte lineage NSCs/TAPs amounted to approximately 7% of all NSCs and TAPs, which is in line with earlier retroviral-fate mapping observations describing a similar oligodendrocyte/neuron output ratio between from the adult SVZ (Menn et al., 2006). Comparison of single cells at a defined stage of the two lineages showed only a modest degree of separation in tSNE plots. While this implies close similarity between the early steps of oligodendroglial and neuronal lineage progression, key differences were associated with specific classes of genes, such as transcription factors. Notably, more closely focusing on lineage-specific features by GO “Protein Class” and “Pathway” analyses revealed that from the earliest quiescent NSC stages to TAPs, the terms related to “transcriptional cues” accounted for most of the differences between the two lineages. In agreement with the findings of the present study, these broad classes of genes have recently been described as enriched in oligodendroglial clusters of adult NSCs/TAPs (Mizrak et al., 2019), similar to observations made during postnatal development in pre-OPC (TAPs) populations (Marques et al., 2018), and elsewhere during embryonic oligodendrogenesis (Klum et al., 2018). Our comprehensive side-by-side comparison of protein classes differentially expressed by the two SVZ lineages unequivocally demonstrates transcriptional control as a major determinant of fate specification and lineage progression and warranted further in-depth characterization of the GRN controlling oligodendrogenesis (see below). Pathway analysis of the most immature stages (qNSCI to TAP) in the two lineages suggested previously unappreciated mechanisms controlling the generation of adult oligodendrocytes from the SVZ. Important pathways detected in the qNSCI/II and pNSC stages of the oligodendrocyte lineage included Wnt signaling, angiogenesis and Notch signaling, all known as important pathways regulating adult oligodendrogenesis (reviewed in (El Waly et al., 2014)). Other key pathway terms derived from this analysis include “Circadian clock”, which has not previously been described as a regulator of oligodendrogenesis, although an earlier study proposed this pathway as a cell-intrinsic timer for inhibiting cell division in postnatal OPCs (Gao, Durand, & Raff, 1997). Interestingly, PDGF signaling was uniquely enriched in OLpNSCs in our analysis and we predict that this oligodendrocyte lineage stage comprises the SVZ cell population previously reported to undergo *in vivo* expansion upon exposure to infused PDGF-A (Jackson et al., 2006; Moore, Bain, Loh, & Levison, 2014). Pathways detected in aNSC and TAP populations comprise “Cell cycle”, “DNA/nucleotide synthesis” and “Transcriptional machineries”, consistent with single cell profiling studies of adult neurogenesis (Dulken et al., 2019; Llorens-Bobadilla et al., 2015). We also identified Notch signaling as a regulator of oligodendrocyte-fated aNSCs and TAPs, which is in agreement with previous observations in the context of early SVZ-derived gliomagenesis (Giachino et al., 2015) and during OPC generation from embryonic NPCs (Cui et al., 2004).

Interestingly, scoring the early oligodendroglial and neuronal lineage cells for their proliferative state revealed instead broad similarities between corresponding cell stages, suggesting that lineage commitment and proliferative expansion are at least in part independently controlled. These similarities were fewer at post-TAP stages, where OPCs and the very earliest neuroblasts (before emigration via the rostral migratory stream) comprised mostly of quiescent cells and proliferative states, respectively. The latter stage, upon pharmacogenomically instructed manipulation for directing specific cell fates in older adult mice, results in its expansion and subsequent dorsal-SVZ-derived OPCs (Azim et al., 2017). In support of these findings, “Nucleic acid binding” and “transcription factor” pathways were most prominent during S phase compared to G1 and G2M phases, suggesting transcriptional modulation associated with access to *de novo* synthesized DNA (e.g. propagation or passive loss of modifications on newly added histones) (Figure 3e). Thus, targeting transcription factor programs regulating cell cycle phases in adult OLTAPs as previously described (Azim et al., 2017), presents additional avenues in promoting the generating of adult-born OPCs.

Establishment of a pan-oligodendrocyte GRN allowed us to investigate transcriptional network organization during lineage progression, defining stage-specific GRN states and their core regulators, which add to well-established broad-oligodendrocyte transcription factors such as *Olig2* and *Sox10*. These transcription factors are well-characterized as binding to the promoters of a large number of genes regulating several aspects of NSC/progenitor behaviors. *Olig2*, aside from regulating transcriptional programs associated with lineage progression and myelination in the post-OPC stages, also represses proneuronal and quiescence genes and activates genes that promote cell cycle entry and oligodendrogenesis in NSCs/NPs (Mateo et al., 2015). A number of *Sox10* target genes overlap with those of *Olig2* in later stage oligodendrocytes (Cantone et al., 2019), while comparatively little is known about oligodendrocyte transcription factor genome occupancy patterns in SVZ-NSCs. In our analysis, *Sox10* mRNA was detected already in OLaNSCs, consistent with its expression downstream of *Olig2* (Liu et al., 2007) and we predict *Sox10* to be downregulating genes expressed in the neuronal lineage, including *Sufu* that represses signaling pathways (e.g. Wnt and Shh signaling) which limit the specification of OPCs (Pozniak et al., 2010).

Surprisingly, transcriptional control of lineage progression was most marked during the actively proliferating oligodendrocyte lineage stages (OLaNSC and OLTAP) and at the OLqNSCII stage, which we explored for predictive functions of transcription factors. In our reconstructed GRN, analyzing transcription factor-target gene interactions in the oligodendrocyte lineage aside from the well-characterized transcriptional regulators, we uncovered novel master regulators. We used our reconstructed GRN for predicting gene regulatory functions of these putative master regulators. Amongst those identified were *Cebpa*, which has been described to regulate genes expressed in the cycling population of the oligodendrocyte lineage (Calella et al., 2007), and *Mitf*, which activates the transcription of the *Dct* gene (Jiao et al., 2006), previously identified as highly expressed in gliogenic SVZ-NSCs (Azim et al., 2015). Transcription factors expressed by OLaNSCs tended to positively regulate TAP-specific genes, thus possibly promoting rapid lineage progression, while the set of TAP-expressed transcription factors suggests concomitant reinforcement of the TAP-specific gene-expression program and repression of differentiation-associated genes (Figure 6e,f). Indeed, target gene repression emerged as more widespread than in the neuronal lineage and may reflect the need for stabilization of an intermediate undifferentiated stage after initial lineage progression and expansion downstream of NSCs (Menn et al., 2006).

Our work identified transcription factors/transcriptional networks in the earlier stages of the oligodendrocyte lineage that had been poorly characterized, and future mechanistic studies will aim to confirm their predicted gene regulatory control of oligodendrogenesis. Finally, given the predicted relevance of environmental signaling and transcriptional control over oligodendrocyte lineage progression, we investigated the ability of extracellular signaling events to modulate core transcriptional regulators of the oligodendrocyte GRN. Several well-established signaling pathways were predicted to stage-specifically impinge on core GRN components, suggesting a measure of environmental control over lineage progression. Thus, while SVZ-derived oligodendrocyte lineages are capable of remarkable intrinsic control over their maturation (F. Ortega et al., 2013), a series of known and novel environmental factors can confer flexibility to this process by regulating the expression of transcriptional modulators of oligodendrocyte lineage progression.

## AUTHORS’ CONTRIBUTIONS

KA was responsible for the conceptualization, data curation, formal analysis, funding acquisition, investigation, methodology, project administration, supervision. FC contributed to writing, validation and data analysis. MC carried out data curation, analysis, investigation validation, and methodology. RA contributed to writing, data curation, formal analysis, investigation, methodology. JV was responsible for funding acquisition, project administration, supervision; HPH: funding acquisition, supervision and project administration. OB was involved in data curation, acquisition and methodology. AB was responsible for writing, methodology, supervision and validation. HPH was responsible for funding acquisition and project administration. PK for funding acquisition, investigation, methodology, project administration, supervision, validation and writing.

## OPEN RESEARCH

Scripts developed for the first time, Cytoscape files, cluster specific gene lists, gene matrices and any other raw data of this study are placed in the repository Github https://github.com/kasumaz/AdultOLgenesis. Assistance for TETRAMER usage can also be requested from Dr Marco A. Mendoza. A user friendly web interface that allows investigators to examine gene expression across all single cells assembled will be made online upon acceptance with the of the lab of Julio ww.curatopes.com/melanoma/). Vera (see for example: https://www.curatopes.com/melanoma/).

## ACKNOWLEDGEMENTS

We wish to thank Dominic Grün at the Max Planck Institute of Immunobiology and Epigenetics in Freiburg, DE for his valuable expertise with the pseudotime analysis. We also wish to thank Dr Marco Antonio Mendoza at Genoscope, FR, for establishment, guidance and optimization of TETRAMER.

## List of abbreviations

aNSC: activated neural stem cell; CNS: central nervous system; DEGs: differentially expressed genes; GO: gene ontology GRN: gene regulatory network; MOL: mature oligodendrocyte; NB: neuroblast; NSC: neural stem cell; oligodendrocyte: oligodendrocyte; OLaNSC: oligodendroglial activated neural stem cell; OLpNSC: oligodendroglial primed neural stem cell; OLqNSCI: oligodendroglial quiescent neural stem cell (subtype I); OLqNSCI: oligodendroglial quiescent neural stem cell (subtype II); OLTAP: oligodendroglial transiently amplifying progenitor; OPC: oligodendrocyte precursor cell; PANTHER: protein annotation through evolutionary relationship; PCA: principal component analysis; pNSC: primed neural stem cell; qNSCI: quiescent neural stem cell (subtype I); qNSCII: quiescent neural stem cell (subtype II); SVZ: subventricular zone; TAP: transiently amplifying progenitor; TETRAMER: TEmporal TRAnscription regulation ModellER TF: transcription factor; TG: target gene; tSNE: t-Distributed Stochastic Neighbor Embedding.

## Supporting Information

**Figure S1:**
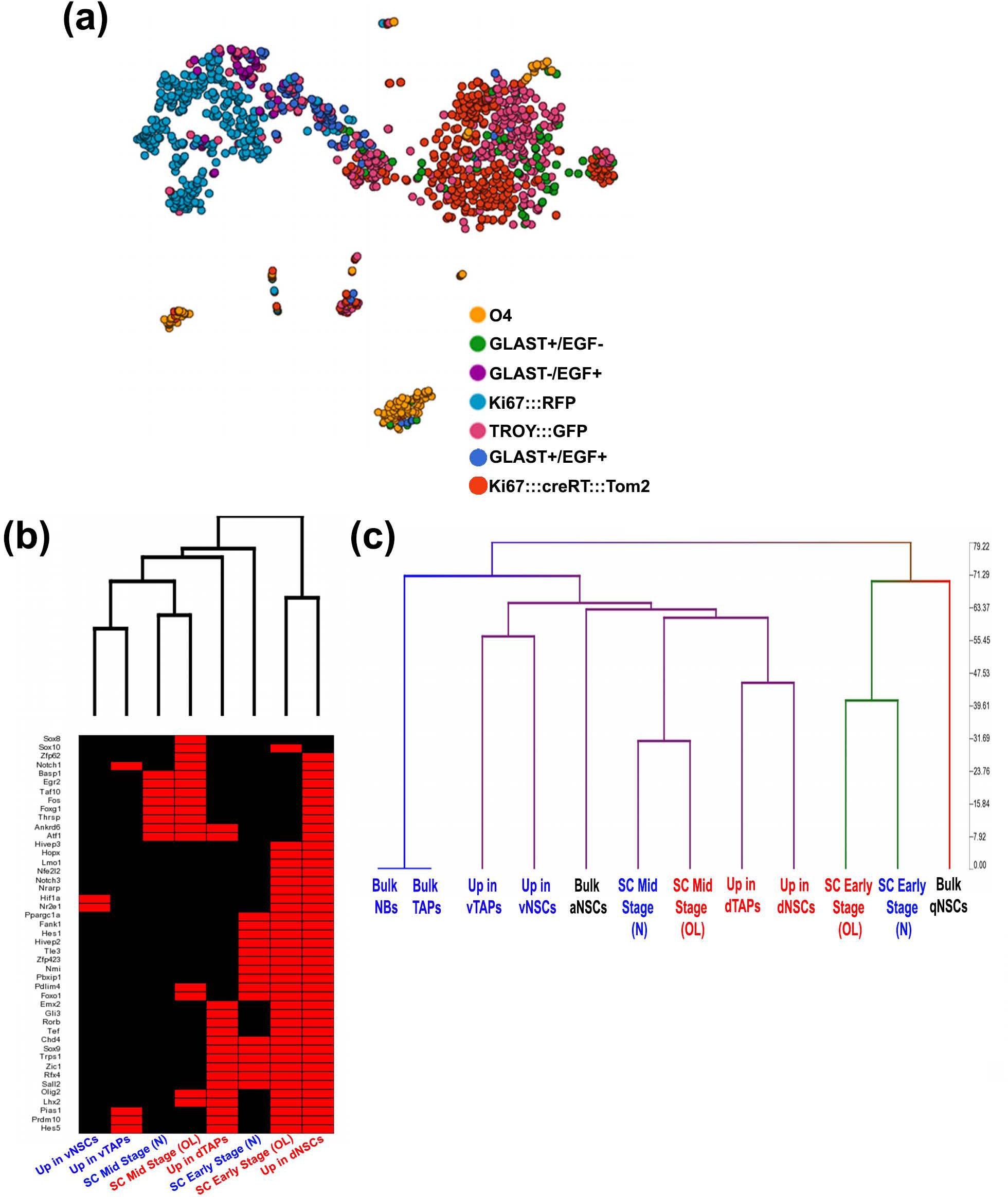
Summary of the datasets processed and overview of the transcription factor expression comparisons with postnatal bulk and adult single analysis. (a) Datasets re-processed illustrating the transgenic reporter and immunofluorescence strategy for isolating cells derived from the adult SVZ. Processed cells are shown as a tSNE plot. (b) Heatmap demonstrating that transcription factors expressed in single cells of OLqNSCI, OLqNSCII and OLpNSC (single cell early stage oligodendrocytes) group closely with postnatal dorsal bulk datasets. Lists of differentially expressed transcription factor-encoding genes from the indicated analyses were rendered as lists of binary values to allow grouping. (c) Clustergram illustrating transcriptomic differences and similarities of datasets generated from bulk versus single cell sequencing. Abbreviations: aNSC: activated neural stem cell; dTAPs: dorsal TAPs; dSVZ: dorsal subventricular zone; N: neuronal; NB: neuroblast; NSC: neural stem cell; oligodendrocyte: oligodendrocyte; OL: oligodendroglial; OPC: oligodendrocyte precursor cell; PANTHER: protein annotation through evolutionary relationship; PCA: principal component analysis; qNSC: quiescent neural stem cells; TAP: transiently amplifying progenitor.

**Figure S2:**
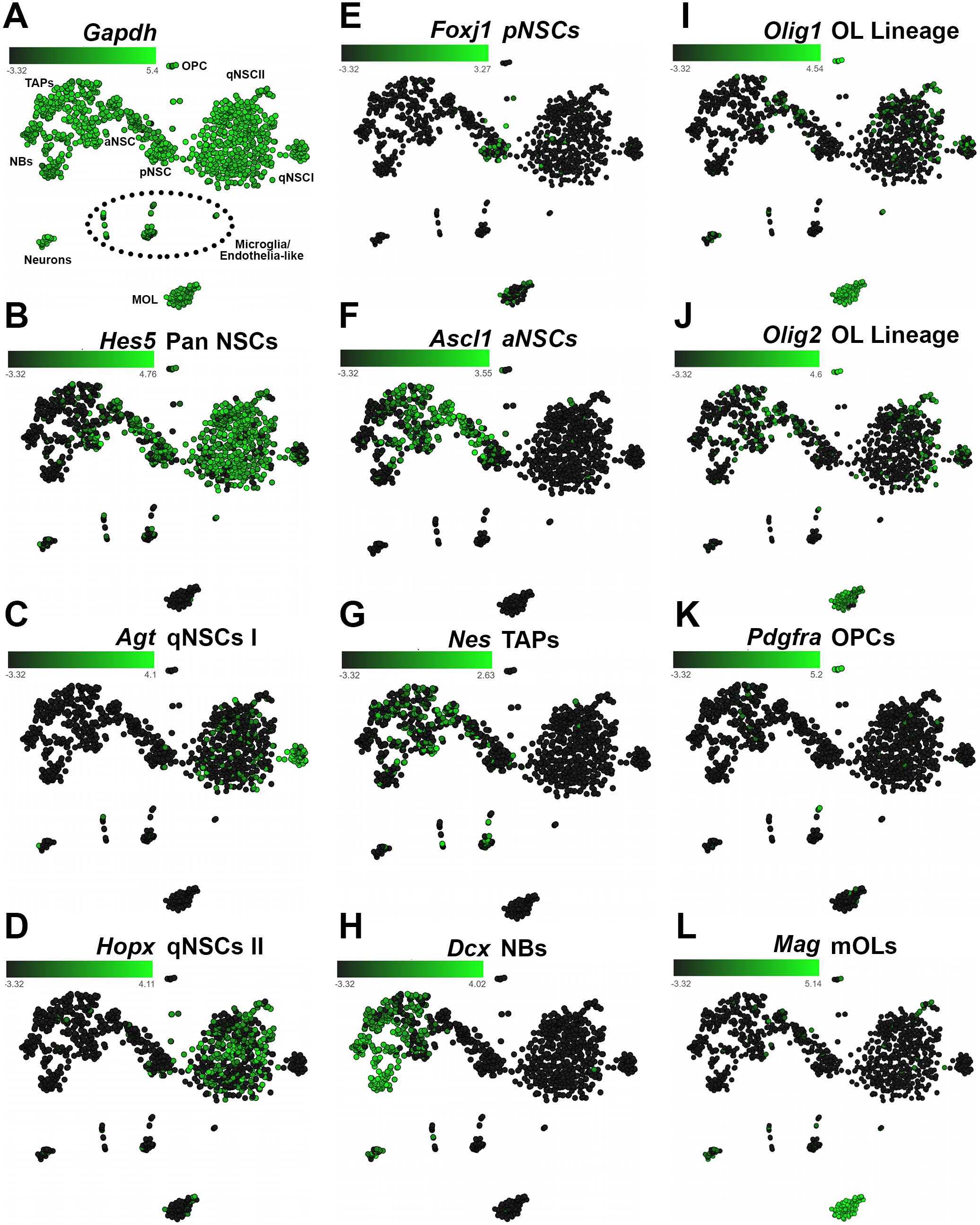
Stage and cell specific marker expression across single cells in tSNE plots. (a) Constant *Gapdh* expression levels across cell types studied. (b-g) Selected markers of stage-specific markers, including pan-early NSC populations (*Hes5* qNSC-pNSC) and neuroblasts (*Dcx*). (i-l) Landmark markers of oligodendrocytes. Abbreviations: mOL: mature oligodendrocyte; NSC: neural stem cell; OPC: oligodendrocyte precursor cell; pNSC: primed neural stem cell; qNSCI: quiescent neural stem cell (subtype I); qNSCII: quiescent neural stem cell (subtype II);TAP: transiently amplifying progenitor.

**Figure S3:**
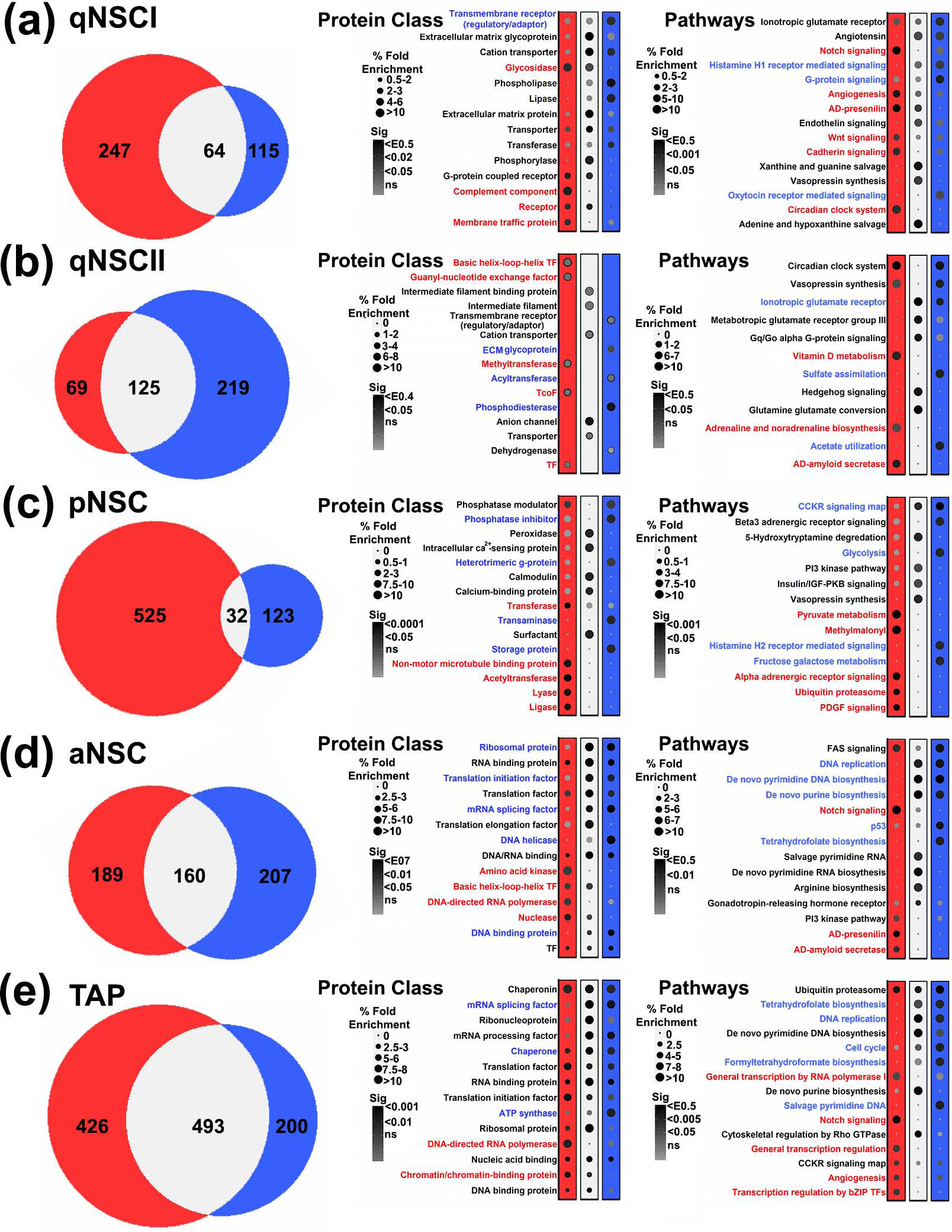
Stage-specific comparisons of oligodendrocyte lineage versus neuronal lineage for PANTHER Protein Class and Pathways. (a-e) Cluster enriched genes for the oligodendroglial (red throughout) and neuronal linage (blue throughout) for each stages analysed were compared for Panther Protein Class and Pathways. The overlapping genes (grey throughout) in each stage were included in the analysis. Ontology terms enriched in the oligodendrocyte lineage, neuronal lineage and common to both lineages are highlighted in red, blue and black text, respectively. Abbreviations: NSC: neural stem cell; pNSC: primed neural stem cell; qNSCI: quiescent neural stem cell (subtype I); qNSCII: quiescent neural stem cell (subtype II);TAP: transiently amplifying progenitor; TF: transcription factor.

**Figure S4:**
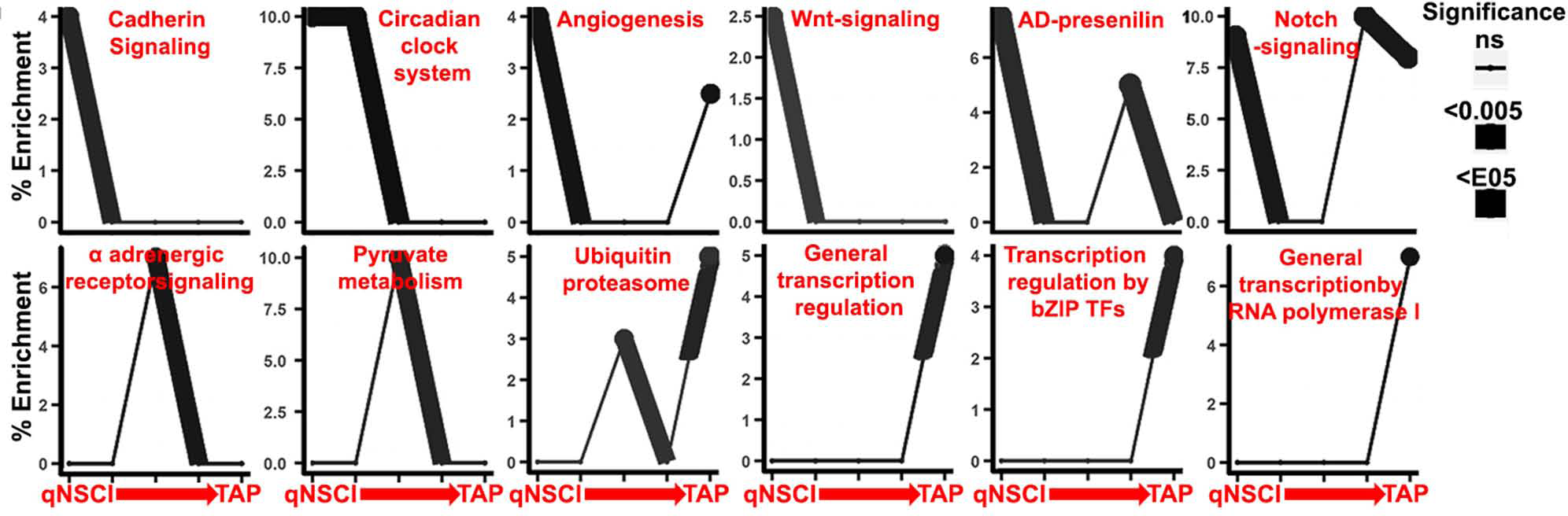
Temporal PANTHER Pathway enrichment in the oligodendrocyte lineage. A time course of selected pathways enriched in the oligodendrocyte lineage from the PANTHER analysis shows enrichment of mechanisms during the course of oligodendrogenesis. Red, grey and blue colours in Venn diagrams signify, oligodendroglial, intersect (common), and neuronal lineage cells, respectively. A few pathways such as Angiogenesis and Notch signalling for example show enrichment in defined stages. Terms have been shorted/abbreviated to fit.

**Figure S5:**
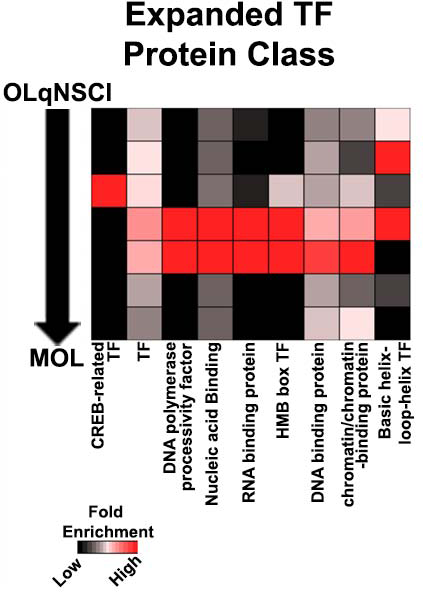
Temporal Transcriptional Regulation in the oligodendrocyte lineage. Protein class terms abundant in the 3 stages in Figure S4 are expanded further as a heatmap for representing modes of transcription factors regulating the oligodendrocyte lineage. TF: transcription factor.

**Figure S6:**
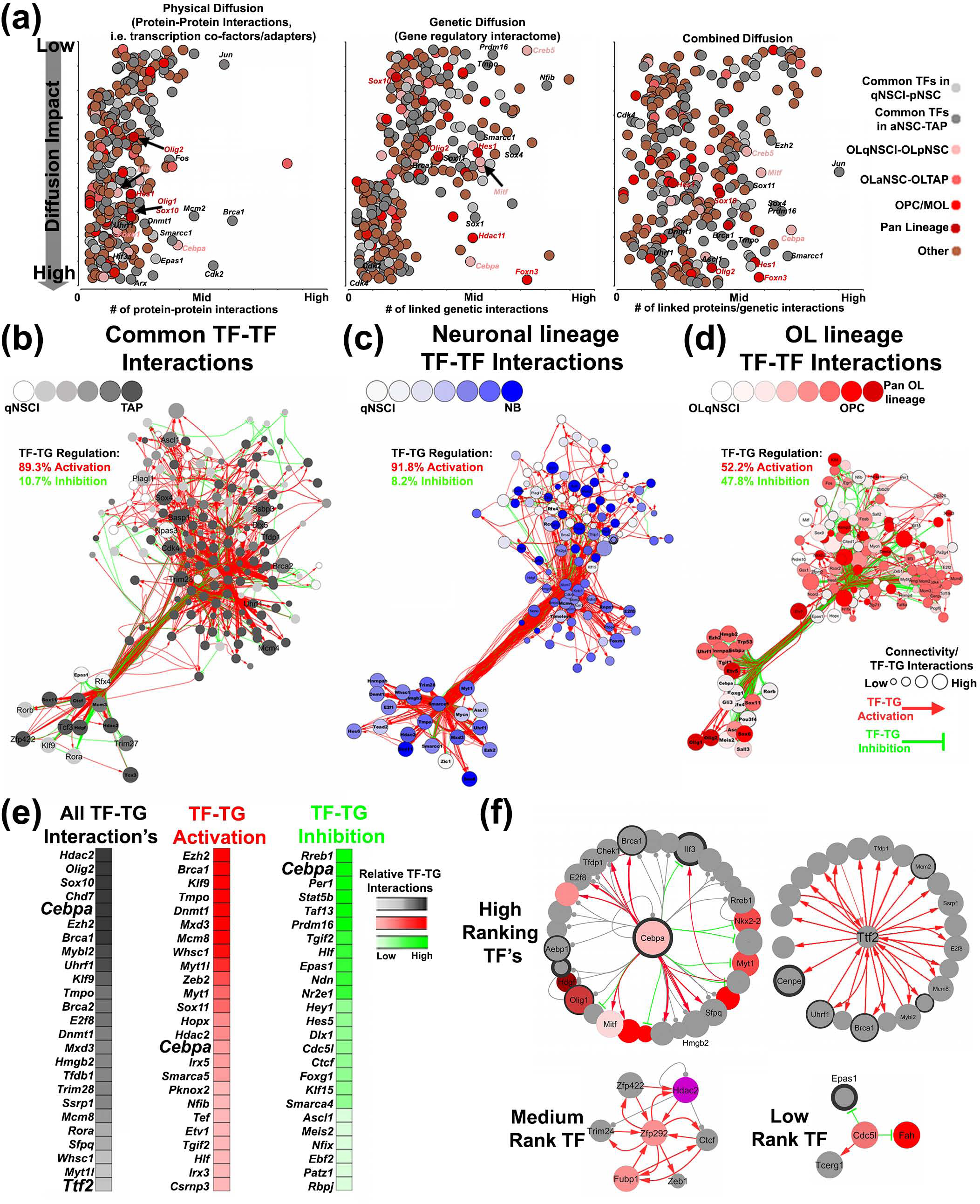
Transcription factor expression in oligodendroglia points to shifts in gene regulatory network states as major drivers of differentiation. (a) Transcription factors expressed in the oligodendrocyte lineage (light red from OLqNSCI to dark red for mature oligodendrocytes) and expressed in both lineages, i.e. commonly expressed transcription factors between the 2 lineages (light grey from qNSCI to dark grey for TAPs) are ranked according to their heat diffusion impact for downstream genetic interactions, physical interactions with other transcription factors and combined. (b-d) Using gene lists for the defined stages for transcription factor expression, those transcription factors expressed in both lineages from qNSCI to TAP stage (b), enriched in the neuronal lineage from qNSCI to the NB stage e and oligodendroglial lineage from OLqNSCI to OPC stage (d). The size of each transcription factor node in the network is relative to its transcription factor-TG (target gene) connectivity index. The numbers of activating (red) or inhibiting (green) interactions from each network are shown as a percentage. (e) The top 25 transcription factors with the highest connectivity index are displayed as heatmaps for the numbers of regulatory interactions with TGs. Heatmaps show darker to lighter colour relative to the numbers of genes regulated by each transcription factor. (f) Examples of higher-, medium-and low-ranking transcription factors. See Figure 4d for description of the legend. Abbreviations: MOL: mature oligodendrocyte; NB: neuroblast; NSC: neural stem cell; OL: oligodendrocyte; OLaNSC: oligodendroglial activated neural stem cell; OLpNSC: oligodendroglial primed neural stem cell; OLqNSCI: oligodendroglial quiescent neural stem cell (subtype I); OLqNSCI: oligodendroglial quiescent neural stem cell (subtype II); OLTAP: oligodendroglial transiently amplifying progenitor; OPC: oligodendrocyte precursor cell; pNSC: primed neural stem cell; qNSCI: quiescent neural stem cell (subtype I); TAP: transiently amplifying progenitor; transcription factor; TG: target gene.

**Figure S7:**
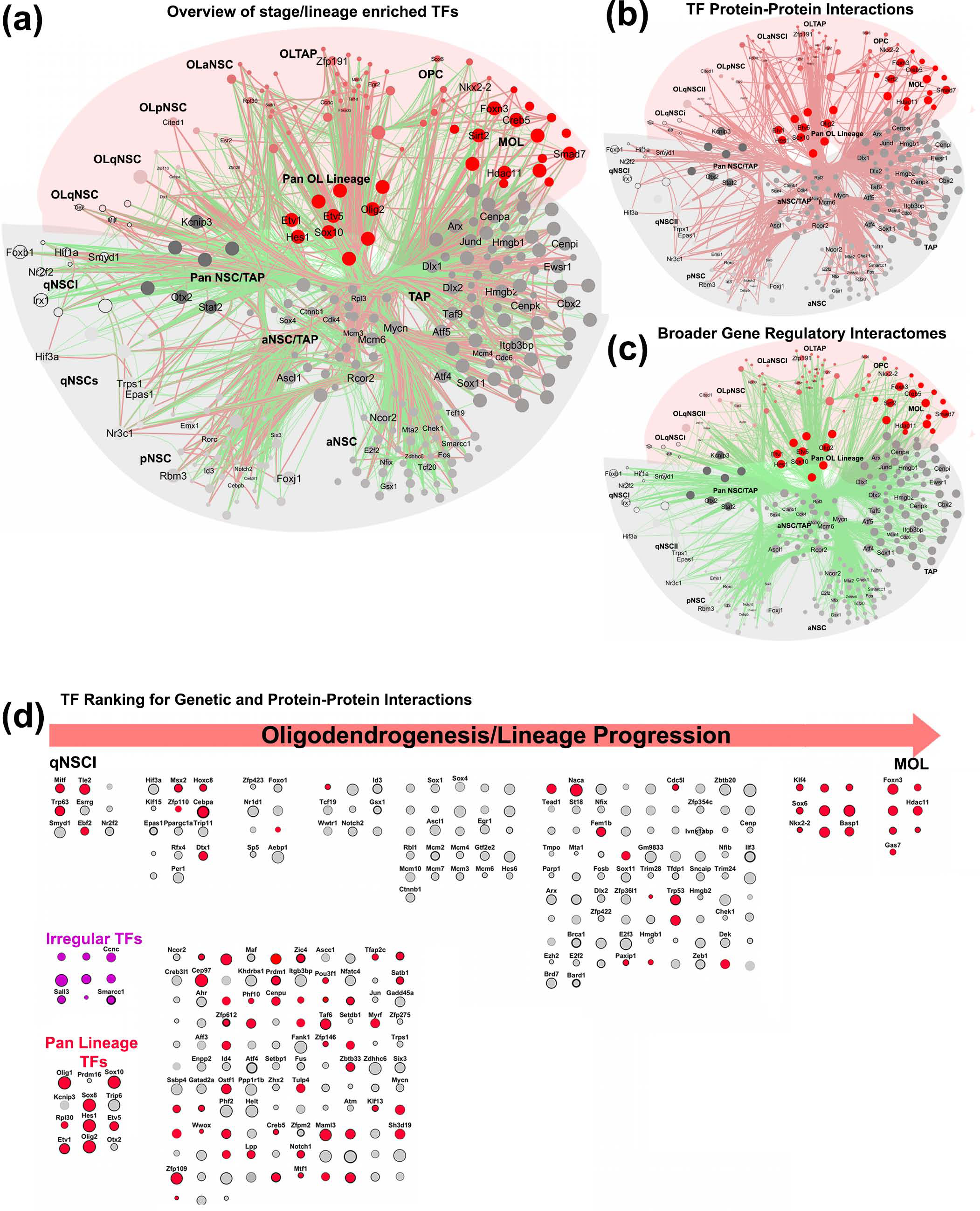
Diffusion ranking for transcription factor functional relationships. (a) Transcription factors expressed in the oligodendrocyte lineage from OLqNSC to MOL stage and those that are common to both oligodendroglial and neuronal lineages were first converted to human gene symbols for maximising the data gathered (information on human gene orthologs are more numerous). (b,c) GeneMania was used to gather functional information on genetic and physical interactions (protein-protein interactions) for the genes sampled. All genes were initially assembled on a network without prior arrangement and the heat diffusion algorithm applied separately for genetic and physical interactions. Node sizes in the networks reflect diffusion rankings for the two combined parameters. Network organized using the CoSE algorithm for force directing nodes within the defined stages. (d) Grading of transcription factors within each stage. Larger to smaller node sizes are relative to the diffusion ranking scores of combined genetic and physical diffusion. Border thickness of each transcription factor directly proportional to protein-protein interaction number. Transcription factors labels are shown for those prioritised and additional transcription factors shown are expressed in the oligodendrocyte lineage with significant expression, whilst the remaining are derived from more stringent criteria. Abbreviations: MOL: mature oligodendrocyte; NSC: neural stem cell; OL: oligodendrocyte: OLaNSC: oligodendroglial activated neural stem cell; OLpNSC: oligodendroglial primed neural stem cell; OLqNSCI: oligodendroglial quiescent neural stem cell (subtype I); OLqNSCI: oligodendroglial quiescent neural stem cell (subtype II); OLTAP: oligodendroglial transiently amplifying progenitor; OPC: oligodendrocyte precursor cell; pNSC: primed neural stem cell; qNSCI: quiescent neural stem cell (subtype I); TAP: transiently amplifying progenitor; transcription factor.

